# A tumor metabolism-angiogenesis-immune axis governs immunotherapy responses

**DOI:** 10.64898/2026.02.23.707524

**Authors:** Inna Serganova, Giorgia Colombo, Francesco Ballesio, Jee Hye Kang, Triantafyllia Karakousi, Tullio V. F. Esposito, Ellen Ackerstaff, Anthony Santella, Ronald Blasberg, Naga Vara Kishore Pillarsetty, Ashley Schreier, Eleni Andreopoulou, Sandra Demaria, Amanda W. Lund, Pier Federico Gherardini, Roberta Zappasodi

## Abstract

Despite the clinical success of immunotherapy, long-lasting benefit remains restricted to a subset of patients. Tumor metabolic adaptation is emerging as a key factor limiting immunotherapy efficacy. We previously found that glycolysis-low tumor variants, compared to the parental glycolytic tumors, better respond to neoadjuvant CTLA-4 immune checkpoint blockade (ICB) therapy. Here, we investigated new rational modalities to restore immune sensitivity of glycolytic tumors by studying how lowering the tumor-cell glycolytic capacity reshapes the tumor microenvironment (TME) to favor long-lasting systemic anti-tumor responses upon immunotherapy.

We found that lowering glycolysis in cancer cells through LDHA-knock down (KD) results in TME displaying normalized vasculature, reduced angiogenic markers, increased high endothelial venules (HEVs), and enhanced recirculation of CD8^+^ T cells both in and out of the tumor. By leveraging public transcriptomic data sets from human solid cancers, we confirmed that glycolysis positively correlates with neo-angiogenesis and inversely correlates with features of vascular normalization and immune cytolytic activity. Moreover, a tumor signature that incorporates glycolysis- and angiogenesis-related genes as positive features, and normal vasculature, HEV, and immune cytolytic activity genes as negative features predicted poor outcomes better than the individual features across most human solid tumor types in the TCGA. To determine the therapeutic implication of these interrelated processes, we asked if targeting the vasculature would restore immunotherapy responses in glycolytic tumors. We found that combining low-dose anti-VEGFR2 with CTLA-4 blockade induces tumor regressions and protection from metastases in glycolytic tumors. These therapeutic effects were associated with vasculature normalization and increased abundance of HEVs and concentrations of lymphangiogenic factors in the TME of glycolytic tumors. Moreover, anti-VEGFR2 with anti-CTLA-4 restored recirculation of anti-tumor CD8^+^ T cells in and out of the TME in glycolytic tumors, with specific increases in intratumoral recruitment and activation of cytolytic CD62L⁺CD44⁺CD8⁺ T cells expressing VEGFR2 and low levels of CTLA-4, suggesting potential novel direct synergistic effects of anti-VEGFR2 and anti-CTLA-4 on CD8^+^ T cells. Conversely, this combination opposed the beneficial immune and vascular TME features of LDHA-KD tumors, indicating tumor-metabolic-dependent effects. Accordingly, we found that standard combined regimens of anti-VEGF and ICB therapy improve survival with respect to ICB alone in patients with glycolysis-high but not glycolysis-low tumors.

Together, these findings indicate that tumor cell glycolysis “primes” the TME for aberrant vascular architecture and T-cell exclusion, and that modulating the tumor vasculature can unravel these mechanisms restoring immune responsiveness. This suggests that tailoring anti-angiogenic and immunotherapy combinations to the tumor glycolytic state and associated vasculature profiles may restore immune surveillance and overcome therapy resistance.

## Introduction

Durable responses to immunotherapy, including immune checkpoint blockade (ICB) and T-cell therapies, occur only in a fraction of patients^1–6^. Immune profiling of exceptional responders to ICB has revealed some key factors associated with cancer regressions, including the presence of tumor-specific T cells able to expand, infiltrate, and functionally persist in the tumor microenvironment (TME)^2,7–10^. The TME of solid tumors poses several obstacles preventing adequate intratumor T-cell infiltration and function, which in turn limit the activity of immunotherapy^11^. One important characteristic of solid tumors is the nutrient-deprived and hypoxic nature of the TME. This is largely due to (1) the metabolic fitness of tumor cells, which rapidly consume available nutrients in the TME to support their high energy demand for rapid proliferation, and (2) the aberrant and immature tumor vasculature that develops as a consequence of dysregulated angiogenesis in the TME.

Cancer cells favor aerobic glycolysis to support their anabolic demands (Warburg effect). Such metabolic adaptation in tumor cells is enabled by the overexpression of glycolytic enzymes – especially lactate dehydrogenase (LDH), which catalyzes the last glycolysis reaction producing lactate^12^. Elevated glycolysis in solid cancers correlates with tumor progression and immune evasion gene signatures^13–15^, as well as poor responses to immunotherapies in patients^16,17^. Moreover, the reported reduced efficacy of immunotherapy in patients with high LDH levels suggests that elevated tumor glycolysis may hamper anti-tumor T-cell function, as observed, for example, in patients with melanoma and triple-negative breast cancer (TNBC)^13,18^.

Vasculature abnormalities in solid tumors constitute another major obstacle toward their effective treatment. Immature and leaky blood vessels in the TME limit drug delivery and T-cell infiltration, while promoting hypoxia, which induces the production of immunosuppressive cytokines, and intratumor accumulation of immunosuppressive cells^19,20^. Moreover, hypoxia leads to a cellular response orchestrated by the hypoxia inducible transcription factor HIF-1, which, among key target genes, up-regulates VEGF-A and ANGPT-2, further promoting neo-angiogenesis^21–23^, as well as LDH-A, hence sustaining glycolysis and lactate production^24^. Importantly, glycolytic cells can directly activate HIF-1 irrespective of low oxygen tension as a consequence of high ATP/ADP ratios that inhibit AMPK^25^ – a kinase that normally phosphorylates and promotes HIF-1α degradation, and also due to elevated lactate concentrations, via metabolic interference (PHD inhibition), redox signaling (ROS), and direct protein modification (lactylation)^22,26,27^. This establishes a positive feedback loop between metabolic adaptation toward glycolysis and neo-angiogenesis.

While the mechanistic relationship between these molecular pathways has been well characterized in a cell-intrinsic manner, how these interrelated tumor cell features shape the TME and response to immunotherapy has not been systematically defined. From a therapeutic perspective, the fact that vasculature abnormalities may be a specific feature of glycolytic tumors suggests that angiogenesis may be a specific vulnerability of these aggressive and immune-refractory tumors. Targeting tumor angiogenesis as an anti-cancer strategy has been extensively investigated and is one of the most common approaches being tested in combination with ICB in patients together with chemotherapy^31,32^. However, the results have been inconsistent so far^28–30^, and we currently lack biomarkers that can guide the optimal use of anti-angiogenic therapies with immunotherapy. Emerging evidence suggests that using anti-angiogenic therapies with the intent of normalizing the tumor vasculature rather than depleting blood vessels may be more efficacious, especially in combination with T-cell based immunotherapies. Recent work in mouse TNBC models^31^ and advanced TNBC patients^31,32^ has shown the therapeutic advantage of low-dose anti-angiogenic therapies (anti-VEGFR2 or apatinib) in combination with PD-1 blockade, which was associated with improved tumor vasculature and T-cell infiltration. However, the precise mechanism leading to tumor vasculature normalization that enhances immunotherapy efficacy remains unclear.

In this study, we investigated the relationship between tumor glycolytic capacity and aberrant tumor vasculature and its role in immunotherapy resistance. Furthermore, we leveraged this information to provide a framework for rationally combining ICB with anti-angiogenic therapies based on the tumor glycolytic state. Our work builds on previous observations that downregulating tumor glycolysis by targeting LDH-A improves CTLA-4 blockade immunotherapy in otherwise immune-refractory solid tumor models^13,33^. Here, we found that dampening tumor glycolysis through selective LDHA knock-down (KD) in cancer cells promotes vasculature remodeling toward its normalization in the TME, which is further amplified by CTLA-4 blockade immunotherapy. By using vasculature-normalizing low-dose anti-VEGFR2 therapy^34^, we demonstrated that improving the tumor vasculature of glycolytic tumors restores their responsiveness to CTLA-4 blockade. Mechanistically, we observed that this combination therapy decreases endothelial cells (ECs) and angiogenic pericytes (PCs), while increasing stabilizing quiescent PCs^35,36^ and high endothelial venules (HEVs)^37^. These vasculature effects were linked to increased infiltration and functionality of intratumor CD8^+^ T cells and greater recirculation of tumor-specific CD8^+^ T cells from the TME to draining lymph nodes (dLNs). These vasculature-driven immune effects of the combination treatment can explain the decrease in circulating tumor cells, and further protection from developing distant metastases after primary tumor resection in the neoadjuvant setting. Notably, these vascular and immune effects were attenuated or reversed by the same combination therapy in glycolysis-defective LDHA-KD tumors, where the vasculature is more functional. In human tumors in the TCGA, we found that the expression of glycolysis-related genes positively correlates with angiogenic vasculature phenotypes, and inversely correlates with HEVs, stabilizing quiescent PCs, and cytolytic activity. Notably, a combined score derived from these tumor features identified patients with unfavorable outcomes, with better predictive power compared to the individual features alone. Moreover, by leveraging a published tumor transcriptomic dataset from patients treated with ICB+/- anti-VEGF^38^, we observed that the combination achieves improved benefit over ICB alone selectively in patients with glycolysis-high tumors.

Taken together, these findings suggest that targeted intervention to normalize the tumor vasculature based on the tumor’s glycolytic state may be a rational and effective approach to improve immunotherapy in aggressive and otherwise immune-refractory solid tumors.

## Results

### Tumor glycolysis correlates with aberrant vasculature features and poor immune function

We previously demonstrated that a neoadjuvant CTLA-4 blockade regimen before resection of primary tumors leads to complete tumor remissions and long-lasting anti-tumor immunological memory if glycolysis is dampened in tumor cells, by using LDHA-KD vs. scramble control (Sc) 4T1 TNBC models^13^. Here, we investigated changes in the stromal TME that could promote systemic anti-tumor immune responses in mice bearing LDHA-KD tumors that were treated with neoadjuvant anti-CTLA-4. By interrogating bulk RNA-sequencing (RNA-seq) data of LDHA-KD *versus* (vs.) Sc 4T1 tumors from mice treated with anti-CTLA-4 or IgG control, we observed major vasculature effects driven by LDHA-KD that were eventually amplified by anti-CTLA-4 in tumors: (1) LDHA-KD tumors displayed significant downregulation of gene signatures for tumor-associated ECs^39^ and angiogenic PCs^35,36^; (2) CTLA-4 blockade tended to promote the expression of gene signatures for quiescent PCs^35^, which stabilize the endothelial walls for correct vasculature function, and maximally upregulated marker genes for HEVs^37,39^ in LDHA-KD tumors (**Suppl. Fig. 1A**). Furthermore, consistent with our prior findings^13^, anti-CTLA-4-treated LDHA-KD tumors displayed the greatest immune cytolytic score, based on *PRF1* (encoding perforin-1) and *GZMA* (encoding granzyme A)^40^ (**Suppl. Fig. 1B**).

To corroborate in the human setting the relevance of these observations in glycolysis-defective LDHA-KD vs. control mouse tumor models, we tested the relationship between glycolysis, aberrant vasculature, and immune dysfunction in the human TCGA data set^41^. In addition to human breast cancer, we extended this analysis to human melanoma and lung adenocarcinoma, as these represent additional tumor types where immunotherapy is commonly used. Across these data sets, we found that expression of the glycolysis gene signature or LDHA alone positively correlated with tumor vasculature and angiogenic PC gene signatures (neo-angiogenic features) and inversely correlated with quiescent PC and tumor HEV gene signatures, and cytolytic activity score (for vasculature normalization and immune infiltration and function, respectively) (**Fig. 1A**; **Suppl. Table 1**). The strength of these correlations was greater in the breast cancer and lung adenocarcinoma data sets that include a higher number of cases (**Table 1**). Overall, this mirrored our results in mice, showing that the tumor vasculature is most aberrant in glycolytic tumors, suggesting conserved mechanisms between mice and humans.

**Figure 1:**
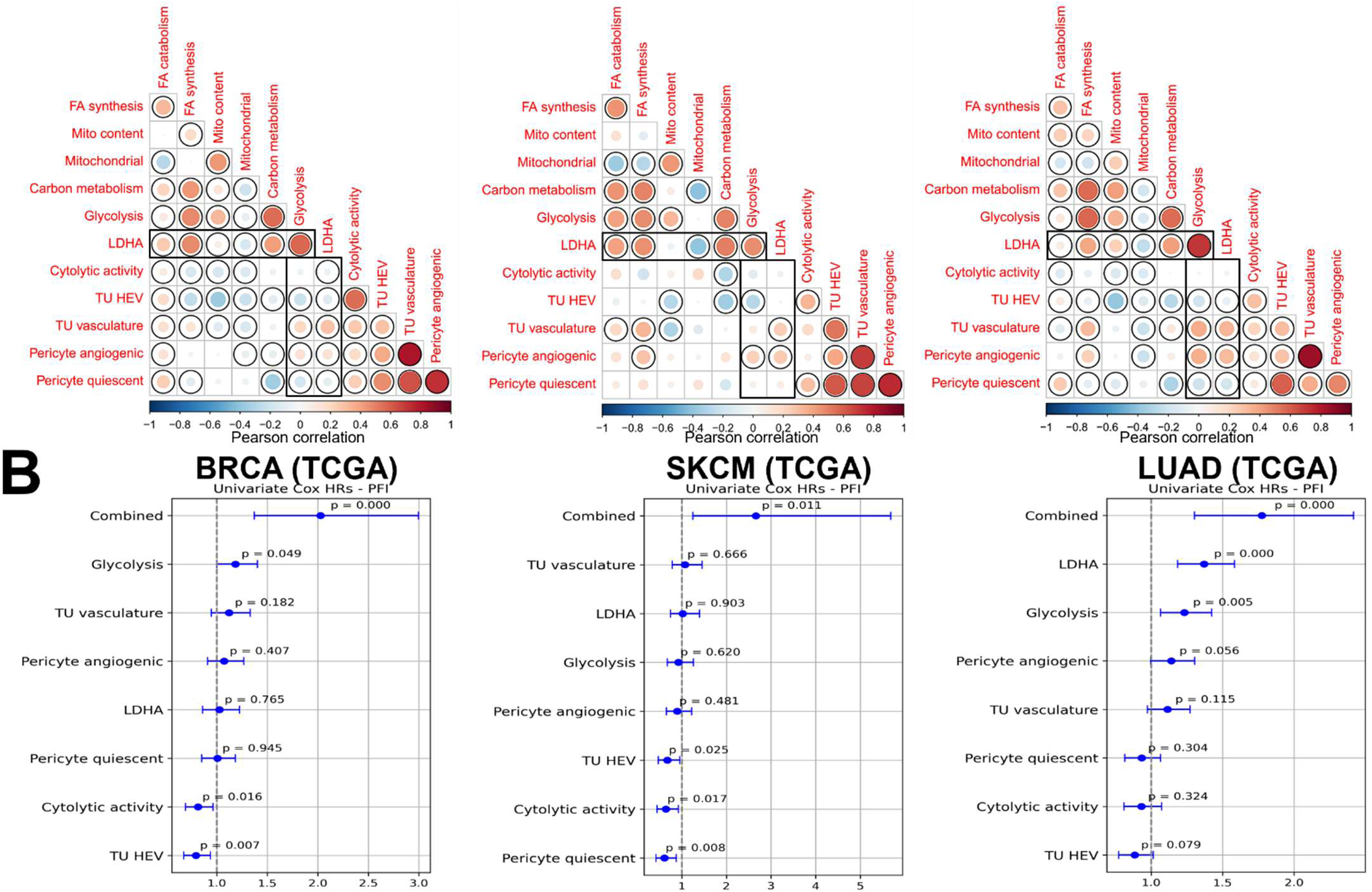
Tumor glycolysis, angiogenesis, and immune evasion are interconnected tumor features and together predict poor outcome in patients. (**A**) Pearson correlation between expression of the indicated signatures (Suppl. Table 1) in the indicated TCGA data sets (black circle, statistical significance). (**B**) HR for a combined signature score incorporating glycolysis, tumor vasculature, angiogenic PC signature scores as positive features and HEV, quiescent PC, and cytolytic signature scores as negative features in comparison with HR for the individual signature scores in primary human breast cancer (BRCA), melanoma (SKMEL), and lung cancer (LUAD) cases from the TCGA data set.

**Table 1:** HR for combined signature score across TCGA solid cancer types in relation to their immunogenicity.

| Dataset | Tumor Type | HR | pvalue | n | TMB (median) | IFNG (median) |
| --- | --- | --- | --- | --- | --- | --- |
| TCGA-LIHC | Liver Hepatocellular Carcinoma | 2.282 | <0.001 | 424 | 2.059 | -4.035 |
| TCGA-ACC | Adrenocortical Carcinoma | 3.482 | <0.001 | 79 | 2.059 | -9.966 |
| TCGA-BRCA | Breast cancer | 2.026 | <0.001 | 1226 | 0.971 | -2.932 |
| TCGA-LUAD | Lung adenocarcinoma | 1.775 | <0.001 | 589 | 4.794 | -1.283 |
| TCGA-HNSC | Head and Neck Squamous Cell Carcinoma | 1.74 | 0.001 | 566 | 2.559 | -1.639 |
| TCGA-KIRP | Kidney Renal Papillary Cell Carcinoma | 3.969 | 0.001 | 323 | 1.529 | -5.012 |
| TCGA-MESO | Mesothelioma | 2.805 | 0.001 | 87 | 0.647 | -1.911 |
| TCGA-LGG | Brain Lower Grade Glioma | 1.89 | 0.001 | 534 | 0.706 | -9.966 |
| TCGA-KICH | Kidney Chromophobe | 8.576 | 0.003 | 91 | 1.353 | -5.574 |
| TCGA-PAAD | Pancreatic Adenocarcinoma | 1.901 | 0.005 | 183 | 0.882 | -3.626 |
| TCGA-SARC | Sarcoma | 1.765 | 0.006 | 265 | 0.971 | -3.308 |
| TCGA-SKCM | Melanoma | 2.662 | 0.011 | 473 | 7.676 | -1.595 |
| TCGA-BLCA | Bladder Cancer | 1.724 | 0.015 | 428 | 4.235 | -3.308 |
| TCGA-CESC | Cervical Squamous Cell Carcinoma and Endocervical Adenocarcinoma | 1.957 | 0.017 | 309 | 2.147 | -0.834 |
| TCGA-COAD | Colon Adenocarcinoma | 1.803 | 0.03 | 514 | 2.824 | -2.681 |
| TCGA-THCA | Thyroid cancer | 2.111 | 0.041 | 572 | 0.206 | -3.383 |
| TCGA-CHOL | Cholangiocarcinoma | 2.685 | 0.064 | 44 | 0.912 | -3.626 |
| TCGA-PRAD | Prostate Cancer | 1.71 | 0.073 | 554 | 0.618 | -4.035 |
| TCGA-UVM | Uveal melanoma | 2.719 | 0.085 | 80 | 0.294 | -9.966 |
| TCGA-TGCT | Testicular Germ Cell Tumors | 2.021 | 0.1 | 156 | 0.353 | 0.276 |
| TCGA-READ | Rectum adenocarcinoma | 1.859 | 0.153 | 177 | 2.265 | -2.826 |
| TCGA-UCEC | Uterine Corpus Endometrial Carcinoma | 1.388 | 0.168 | 585 | 1.882 | -2.826 |
| TCGA-STAD | Stomach adenocarcinoma | 0.766 | 0.227 | 448 | 2.853 | -1.938 |
| TCGA-PCPG | Pheochromocytoma and Paraganglioma | 1.905 | 0.24 | 187 | 0.206 | -9.966 |
| TCGA-KIRC | Kidney renal clear cell carcinoma | 1.313 | 0.282 | 610 | 1.353 | -1.994 |
| TCGA-ESCA | Esophageal carcinoma | 1.133 | 0.557 | 198 | 2.618 | -2.466 |
| TCGA-OV | Ovarian | 1.095 | 0.582 | 429 | 1.676 | -3.626 |
| TCGA-UCS | Uterine Carcinosarcoma | 1.213 | 0.698 | 57 | 1.265 | -5.574 |
| TCGA-LUSC | Lung squamous cell carcinoma | 1.063 | 0.755 | 552 | 5.676 | -1.355 |
| TCGA-THYM | Thymoma | 0.884 | 0.83 | 122 | 0.206 | -2.727 |
| TCGA-GBM | Glioblastoma | 0.988 | 0.96 | 175 | 1.294 | -9.966 |

TMB, tumor mutational burden. Green-to-red color scale illustrates values from low-to-high in each column.

According to the reported function of HEVs to serve as the main T-cell entry site in the TME^42^, the HEV gene signature was the one among all vasculature features tested that most strongly correlated with intratumor T-cell infiltration and with overall leukocyte infiltration both in our murine TNBC RNAseq data set and human TCGA data (**Suppl. Fig. 1C; Suppl. Table 1**).

### A tumor glycolysis-vasculature-immune signature predicts survival in patients

Given the relationships between tumor glycolysis, vascular phenotypes, and poor immune function in both murine and human tumors (**Fig. 1A**, **Suppl. Fig. 1A**) and the negative impact of these features on immunotherapy outcomes in our mouse model (**Suppl Fig. 1B**), we sought to determine the implications of these interrelated processes in tumor progression in patients. To this end, we generated a combined signature score incorporating glycolysis, tumor vasculature, and angiogenic PC signature scores as positive features and tumor HEV, quiescent PC, and cytolytic activity signature scores as negative features. We scored this combined signature in each tumor sample. We tested its impact on the progression-free interval (PFI) of the corresponding patient within the breast cancer, melanoma, and lung adenocarcinoma TCGA data sets. Across these data sets, we found that the combined signature score, compared to each signature score individually, resulted in the most elevated hazard ratio (HR) with statistical significance (**Fig. 1B**). In contrast, the individual signatures were never significantly predictive of outcome in all 3 data sets. We then calculated the HR of the combined signature across all solid tumor types in the TCGA data set. We found it to be significantly associated with reduced PFI in 16 of 31 histological types, including highly aggressive and immune-cold tumors (e.g. liver, cancer, pancreatic cancer, and mesothelioma; **Table 1**). Importantly, the prediction potential of the combined signature did not correlate with common parameters used to estimate tumor immunogenicity (e.g. tumor mutational burden, TMB; IFNG expression; **Table 1**).

Taken together, these results suggested a potential cause-and-effect relationship between the tumor glycolytic potential and aberrant vasculature impacting anti-tumor immunity and immunotherapy responses.

### Decreasing tumor cell glycolysis improves vasculature and T-cell infiltration in the TME

To validate our transcriptomic findings and mechanistically investigate the link between the tumor glycolytic potential, aberrant vasculature, and T-cell exclusion in the TME, we characterized these features at the cellular level in our established glycolysis-defective LDHA-KD and glycolytic Sc tumor models^13,22^. For this, we used the 4T1 TNBC and B16F10 (hereafter B16) melanoma models, which represent highly glycolytic, immune-excluded, and immunotherapy-resistant tumor models from two distinct murine genetic backgrounds (Balb/c and C57BL/6, respectively). As previously reported^13,22^, LDHA-KD compared to Sc tumor cells displayed decreased glycolytic rates by Seahorse profiles (**Suppl. Fig. 2A**), grew slower in immunocompetent mice (**Fig. 2A; Suppl. Fig. 2B**), and showed greater T-cell infiltration in the TME (**Fig. 2B; Suppl. Fig. 2C**), with similar trends observed in 4T1 and B16 models. In addition, LDHA-KD tumor cells secreted decreased levels of VEGF-A (**Suppl. Fig. 2D**), confirming our prior findings in 4T1^22^ and extending them to the B16 model. We next examined how the vasculature is altered in these tumor models, focusing on the same vessel cell types that showed transcriptomic changes in LDHA-KD 4T1 tumors responding to CTLA-4 blockade (**Suppl. Fig. 1A**). Consistent with those data, time course analysis of 4T1-KD vs. Sc tumors showed reduced frequencies of ECs and a phenotypic switch of the PC populations from angiogenic (NG2^+^PDGFRb^-^) to quiescent (NG2^-^PDGFRb^+^) at the cellular level (**Fig. 2C-E; Suppl. Fig. 2E, F**). In addition, the phenotype of these vascular cell types was deeply affected in LDHA-KD vs. Sc 4T1 tumors, with (1) up-regulation of IFN-γ-responsive molecules MHCs and PD-L1 in both ECs and PCs (**Fig. 2C-E**); (2) EC modulation of molecules supporting T-cell infiltration (ICAM-1 up-regulation for T-cell adhesion; FasL down-regulation, for improved survival of transmigrating effector T cells; up-regulation of sulfate sialomucins ligands for CD62L detected by the MECA-79 antibody which marks HEVs) (**Fig. 2C**); and (3) an overall rebalancing of the full angiogenic program promoting EC expression of vasculature integrity molecules (Tie-2) at the cost of neo-vasculature generation (as indicated by decreases in CD34^+^CD31^+^ progenitor (p)ECs, likely impacting on the overall formation of CD31^+^ ECs) (**Fig. 2C**). We repeated these analyses in the B16 model (C57BL/6 mouse genetic background) by performing readouts at a time point of tumor growth where the tumor size was similar between LDHA-KD and Sc tumors obtaining similar results (**Suppl. Fig. 2B, G-J**). Supporting these vasculature changes, tissue interstitial fluid (TIF) of 4T1-KD vs. Sc tumors contained reduced concentrations of hemangiogenic factors (Ang-2, VEGF-A), and trends towards increased concentrations of T-cell effector cytokines (IL-2, IFN-γ), which were surprisingly paralleled by an increase in lymphangiogenic factors (VEGF-C, VEGF-D) (**Fig. 2F**).

**Figure 2:**
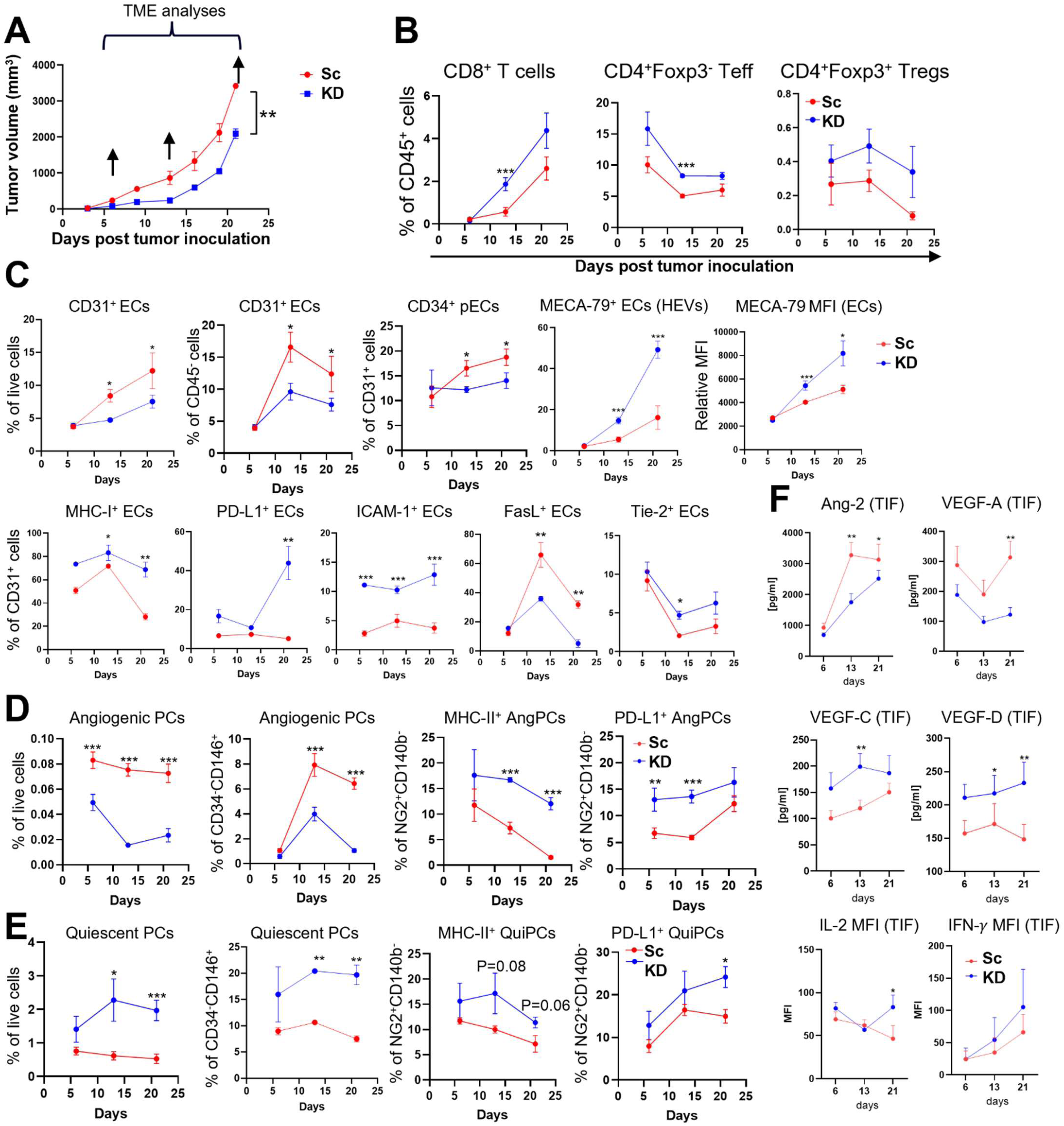
Tumor LDHA-KD reshapes the tumor vasculature phenotype and tumor hem/lymphangiogenic potential. (**A**) Primary tumor growth curves of 4T1 LDHA-KD vs. Sc tumors in WT mice; (**B**) flow cytometry quantification of intratumor T-cell subsets at the indicated time points of tumor growth as in (A); (**C-E**) flow cytometry quantification of tumor-associated ECs (C), angiogenic (Ang) PCs (D), and quiescent (Qui) PCs (E) along with the indicated markers at the indicated time points of tumor growth as in (A); (**F**) Quantification of the indicated soluble angiogenic and immune factors at protein level by Luminex beads-based immunoassay in tissue interstitial fluid (TIF) of tumors at the indicated time points of tumor growth as in (A). Data are mean ± SEM (n=5) of one representative out of 3 independent experiments (multiple unpaired *t* test *, P<0.05; **, P<0.01; ***P<0.001). MFI, median fluorescence intensity.

These findings point to a general mechanistic role of the tumor-cell-intrinsic glycolytic capacity in shaping the tumor vasculature *in vivo,* with potential consequences for the anti-tumor immune response.

### Decreasing tumor cell glycolysis leads to structural vasculature normalization in the TME

In keeping with reduced LDHA levels in cancer cells, LDHA-KD tumors display reduced glucose avidity *in vivo* (as measured by [^18^F]FDG/PET; **Fig. 3A, Suppl. Fig. 3A, B**) and greater glucose abundance in the TME (as measured by spatial MALDI imaging metabolomics^43^; **Fig. 3B**) compared to glycolytic Sc tumor cells. We thus investigated metabolic, structural, and functional changes of the tumor vasculature arising as a possible consequence of these metabolic changes in tissue, using the 4T1 model.

**Figure 3:**
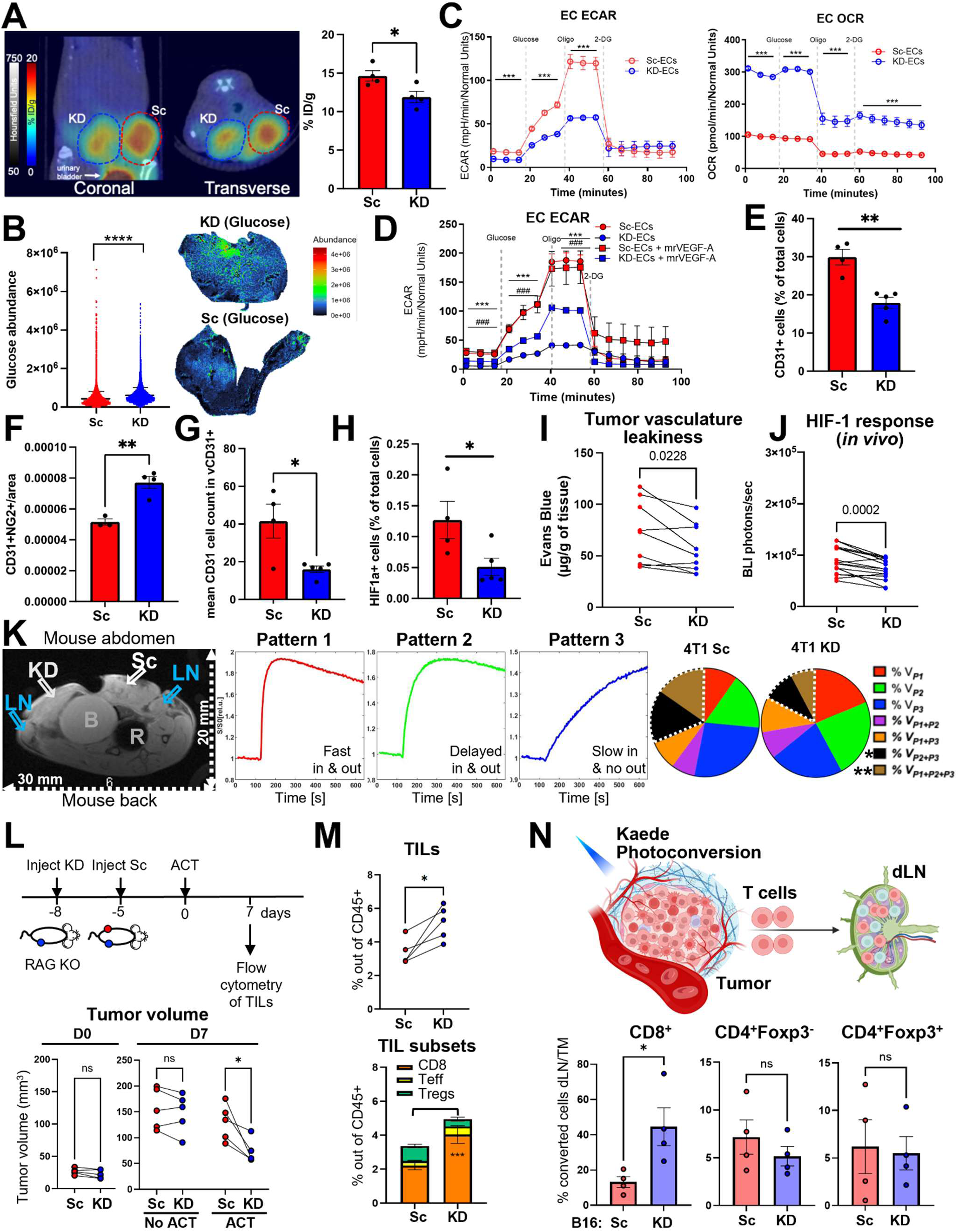
Tumor vasculature normalization in glycolysis-defective vs. glycolytic tumors. (**A**) Representative coronal and transaxial fused PET/CT rendering (PET in colormap, CT in greyscale) and tumor biodistribution of *in vivo* glucose uptake using [^18^F]FDG in mice (n=4) bearing 4T1-KD and 4T1-Sc tumors in contralateral m.f.p. at day 11-14 of tumor growth to equalize tumor volumes. (**B**) Quantification and representative images of pixel intensity for glucose detected by MALDI imaging metabolomics in 4T1-KD and 4T1-Sc tumor tissue. (**C**) Metabolic analysis of EC isolated from Sc and KD tumors by Seahorse, showing extracellular acidification rates (ECAR) and oxygen consumption rates (OCR), and (**D**) Seahorse analysis of Sc- and KD-derived ECs treated with or without murine recombinant (mr)VEGF-A (10 ng/ml). Significance is represented with (*) for control Sc-ECs vs. control KD-ECs, and with (#) for control KD-ECs vs KD-ECs+mrVEGF-A ((n=8/group; Data are mean +/- SD from one representative out of 2-3 independent experiments). (**E-H**) IF quantification of % CD31 ECs (E), fractions of CD31^+^ vessels covered by PCs (F), vasculature complexity by mean CD31^+^ cell numbers in CD31^+^ vessel structures (G), % HIF-1a^+^ cells (H) in 4T1-Sc and KD tumors (n=3-5). (**I, J**) Vessel leakiness by Evans blue assay in 4T1-Sc and 4T1-KD tumors on day 10 and 13 of growth respectively (I), and (J) *in vivo* HIF-1 reporter activity in day 7 vs. day 10 4T1-Sc and 4T1-KD tumors injected 3 days apart in contralateral m.f.p. of WT mice (n=17, 3 experiments combined). (**K**) Representative DCE MR images (showing contralateral 4T1-Sc and 4T1-KD tumors; enlarged dLNs; filled bladder, B; and rectum, R, with inserted rectal temperature probe) (left), and representative average normalized signal-time curves of contrast agent uptake (wash-in and wash-out) behavior of well-vascularized (Pattern 1, typically well-oxygenated), vascularized (Pattern 2, hypoxic) and non-vascularized (Pattern 3, includes necrotic) tumor areas (middle), and (right) related quantification of volume fractions of the indicated tumor perfusion patterns in 4T1-Sc and 4T1-KD tumor tissue (n=12, 2 separate cohorts combined; dotted white line indicates patterns that are significantly decreased in 4T1-KD vs. 4T1-Sc tumors). (**L,M**) Experiment schema with tumor volume (L), and (M) flow cytometry quantification of tumor infiltrating T cells upon adoptive transfer of tumor-specific T cells into RAG2 KO mice bearing contralateral orthotopic 4T1-Sc and 4T1-KD tumors. (**N**) Flow cytometry quantification of proportions of tumor converted T cells (Kaede red^+^) recovered in dLN relative to the same converted T-cell subsets in tumors (TM) of Kaede mice implanted with B16-Sc or B16-KD (n=4/each). Data are mean +/- SE from one representative out of 2-3 independent experiments. 2-sided unpaired (A-H, N) or paired (I-M) *t* test. *, P<0.05; **,P<0.01; ***, P<0.001; ****P<0.0001.

First, we isolated CD31^+^ ECs from LDHA-KD and Sc 4T1 tumors and performed metabolic profiling by Seahorse (**Fig. 3C**) and SCENITH (Single Cell ENergetIc metabolism by Profiling Translation inhibition)^44^ (**Suppl. Fig. 3C**). We found that ECs from glycolysis-low tumors decreased their extracellular acidification rates and increased oxygen consumption rates, implying a shift from a glycolytic phenotype toward OXPHOS-driven energy production (**Fig. 3C**). By SCENITH, we confirmed ECs from glycolysis-low tumors gained in mitochondrial dependence at the cost of losing glycolytic dependence (**Suppl. Fig. 3C**). The direct relationship between tumor cell and intratumoral EC glycolytic capacity suggests that competition for glucose in the TME does not necessarily affect intratumoral EC metabolic activity and that other features of glycolytic tumors may support EC phenotypes, such as for example hemangiogenic factors that are more abundant in glycolytic tumors (e.g. **Fig. 2F, Suppl. Fig. 2D**). Accordingly, we found that *in vitro* culture with recombinant VEGF-A promoted a glycolytic phenotype in ECs derived from LDHA-KD tumors but did not further upregulate glycolysis in ECs from Sc tumors (**Fig. 3D**).

Second, we analyzed vessel structures by immunofluorescence (IF) staining. In keeping with flow cytometry data (**Fig. 2C**), we found reduced frequencies of CD31^+^ ECs in LDHA-KD tumors by IF (**Fig. 3E**), which is also consistent with the decreased glycolytic activity of ECs in these tumors (**Fig. 3C, D; Suppl. Fig. 3C**). As a result, LDHA-KD tumors contained a greater fraction of CD31^+^ vessel structures covered by PCs (**Fig. 3F; Suppl. Fig. 3D**), with vessel structures displaying overall reduced complexity (by the average number of CD31^+^ ECs in these structures) (**Fig. 3G**; **Suppl. Fig. 3D**). These features were associated with a reduced proportion of HIF-1α^+^ cells, suggesting decreased hypoxia in tissue, possibly due to improved vasculature functionality in the TME of glycolysis-defective tumors (**Fig. 3H; Suppl. Fig. 3D**). Similarly, in primary human TNBC cases, we observed decreased vessel structures covered by PCs and increased vessel complexity in tumor tissue compared to normal adjacent tissue (**Suppl. Fig. 3E**).

Third, we performed Evans blue extravasation assays in LDHA-KD and Sc tumors implanted in contralateral mammary fat pad (m.f.p.) of the same mice (to control for main physiology parameters) and found reduced vasculature leakiness in glycolysis-defective (LDHA-KD) vs. glycolytic (Sc) tumors (**Fig. 3I**). Of note, this effect was independent of the presence of an adaptive immune system, as it occurred to the same extent in wild type (WT) and RAG knock out (KO) mice (which lack mature lymphocytes) (**Suppl. Fig. 3F**). In bilateral tumors implanted in WT mice, we also measured HIF-1 activity in cancer cells by using tumor cells transduced with a HIF-1 exGaussia luciferase (exGLuc) reporter system that is detected by *in vivo* bioluminescence imaging (BLI)^22^, showing consistent reductions of this parameter in LDHA-KD vs. Sc tumors (**Fig. 3J**).

Fourth, we analyzed dynamic vessel perfusion in bilateral LDHA-KD and Sc tumors by dynamic contrast-enhanced (DCE) magnetic resonance imaging (MRI) and observed improved patterns of contrast uptake and wash out in LDHA-KD glycolysis-defective tumors (**Fig. 3K; Suppl. Fig. 3G**). Specifically, LDHA-KD tumors displayed reduced proportions of tissue with mixed perfusion/leakage patterns (P2+P3 and P1+P2+P3), with trends toward increased proportions of tissue with improved single perfusion patterns (P1 and P2) (**Fig. 3K; Suppl. Fig. 3G**). These improved vasculature patterns were independent of the primary tumor size, which was equalized by staggering LDHA-KD and Sc tumor implantation (**Suppl Fig. 3H**), whereas they were linked to reduced invasion of the dLNs by LDHA-KD glycolysis-defective vs. glycolytic Sc tumors (**Suppl Fig. 3I**).

Finally, we tested the impact of these vasculature structural differences on T-cell trafficking both in and out of the TME. To test for the former, we adoptively transferred anti-tumor T cells into RAG KO mice bearing contralateral 4T1 LDHA-KD and Sc tumors and compared T-cell recruitment within the two TMEs in each mouse. As 4T1-specific T cells, we used T cells from 4T1 LDHA-KD- and Sc-immunized mice that we activated and expanded with anti-CD3/anti-CD28 before adoptive transfer. Once again, we implanted 4T1 LDHA-KD and Sc tumors 3 days apart so that they reached similar sizes at the time of adoptive transfer (**Fig. 3L**). Importantly, 7 days after transfer, we observed significant reductions in LDHA-KD vs. Sc tumor growth (**Fig. 3L**) coupled with significantly increased T-cell recruitment to LDHA-KD vs. Sc tumors (**Fig. 3M**), indicating improved intratumor infiltration of functional anti-tumor T cells in glycolysis-defective vs. glycolytic tumors. Then, we tested if the improved vasculature in glycolysis-defective tumors could also enable better recirculation of tumor-infiltrating T cells to the periphery, potentially supporting systemic immunosurveillance. To this end, we took advantage of transgenic mice constitutively expressing the photoconvertible fluorescent Kaede protein, which turns from green to red upon exposure to violet light^45^, following a protocol optimized to track leukocyte egress from intradermal tumors to dLN^46^. Because Kaede mice are in the C57BL/6 genetic background, we employed the syngeneic B16 models in these experiments. Twenty-four hours after localized photoconversion of similar-size LDHA-KD and Sc tumor nodules, greater fractions of tumor photoconverted CD8^+^ T cells were recovered in dLNs of mice bearing LDHA-KD vs. Sc B16 tumors (**Fig. 3N**). This indicates more efficient egress of CD8^+^ T cells from glycolysis-defective compared to glycolytic tumors. Other immune cell subsets did not show significant tumor-egress differences between the two tumor metabolic conditions (**Fig. 3N** and not shown). TIL egress to dLN requires transit through lymphatic vessels; however, we did not observe substantial or consistent quantitative changes in lymphatic vessel cells between LDHA-KD and Sc tumors (not shown), potentially indicating differences at the functional level.

Overall, these results further strengthen the causal implication of the tumor-cell glycolytic state in orchestrating the tumor vasculature architecture, and consequent recirculation of T cells in and out of the TME.

### Targeting neo-angiogenesis restores immunotherapy responses in glycolytic tumors

Our findings suggested a mechanism whereby tumor cell glycolytic capacity drives vasculature aberrations in the TME (**Fig. 2**; **Fig. 3A-K; Suppl. Fig. 1A; Suppl. Fig. 2, 3**), in turn affecting T-cell trafficking (**Fig. 3L-N**) and ultimately promoting poor outcomes and resistance to immunotherapy^13^ (**Fig. 1**; **Suppl. Fig. 1B**). To determine the causal implication of aberrant vasculature downstream tumor glycolytic capacity in immunotherapy resistance, we sought to determine whether normalizing the tumor vasculature could restore the response of glycolytic tumors to immunotherapy. To this end, we selected an optimized low-dose regimen of an anti-VEGFR2 blocking antibody with reported vasculature normalization function^34^, given its superior therapeutic and vascular effects compared to VEGF blockade in the 4T1 model in our hands (not shown). We combined this anti-VEGFR2 treatment with a neoadjuvant anti-CTLA-4 regimen, to which we showed glycolytic 4T1 tumors are completely resistant as opposed to glycolysis-defective 4T1 tumors^13^. We found that the combination therapy restored responses against metastatic disease progression from glycolytic 4T1 tumors after surgery, extending survival (**Fig. 4A, B**). The size of primary tumor was not a confounding factor for these effects as it was similar across groups at surgery (**Fig. 4C**). Mechanistically, we found that these therapeutic effects were associated with profound vasculature remodeling in the TME, indicating acquisition of a phenotype closer to the one described for the vasculature of glycolysis-defective tumors (**Fig. 2, 3**). Specifically, vasculature permeability decreased (**Fig. 4D**) along with frequencies of ECs and angiogenic PCs (**Fig. 4E**), and HEVs and inflamed vasculature phenotypes (i.e. PD-L1 expression) increased (**Fig. 4E**) in tumors treated with the combination therapy. Accordingly, the concentration of hemangiogenic factors (Ang-2, VEGF-A) decreased, and the concentration of lymphangiogenic factors (VEGF-C, VEGF-D) along with that of T-cell effector cytokines (i.e. IFN-γ and IL-2) increased in TIF from tumors treated with the combination therapy compared to control or single-agent treatments (**Fig. 4F**). While the amount of total intratumoral CD45^+^ leukocytes did not change significantly, tumor-infiltrating T cells increased upon the combination treatment, with specific expansion of CD4^+^Foxp3^-^effector T cells (Teff) and CD8^+^ T cells, while the frequency of regulatory T cells (Tregs) remained stable (**Fig. 4G**). Notably, most of these effects were most pronounced upon the combination treatment, indicating synergistic vasculature and immune activity through completely independent axes targeted by anti-VEGFR2 and anti-CTLA-4.

**Figure 4:**
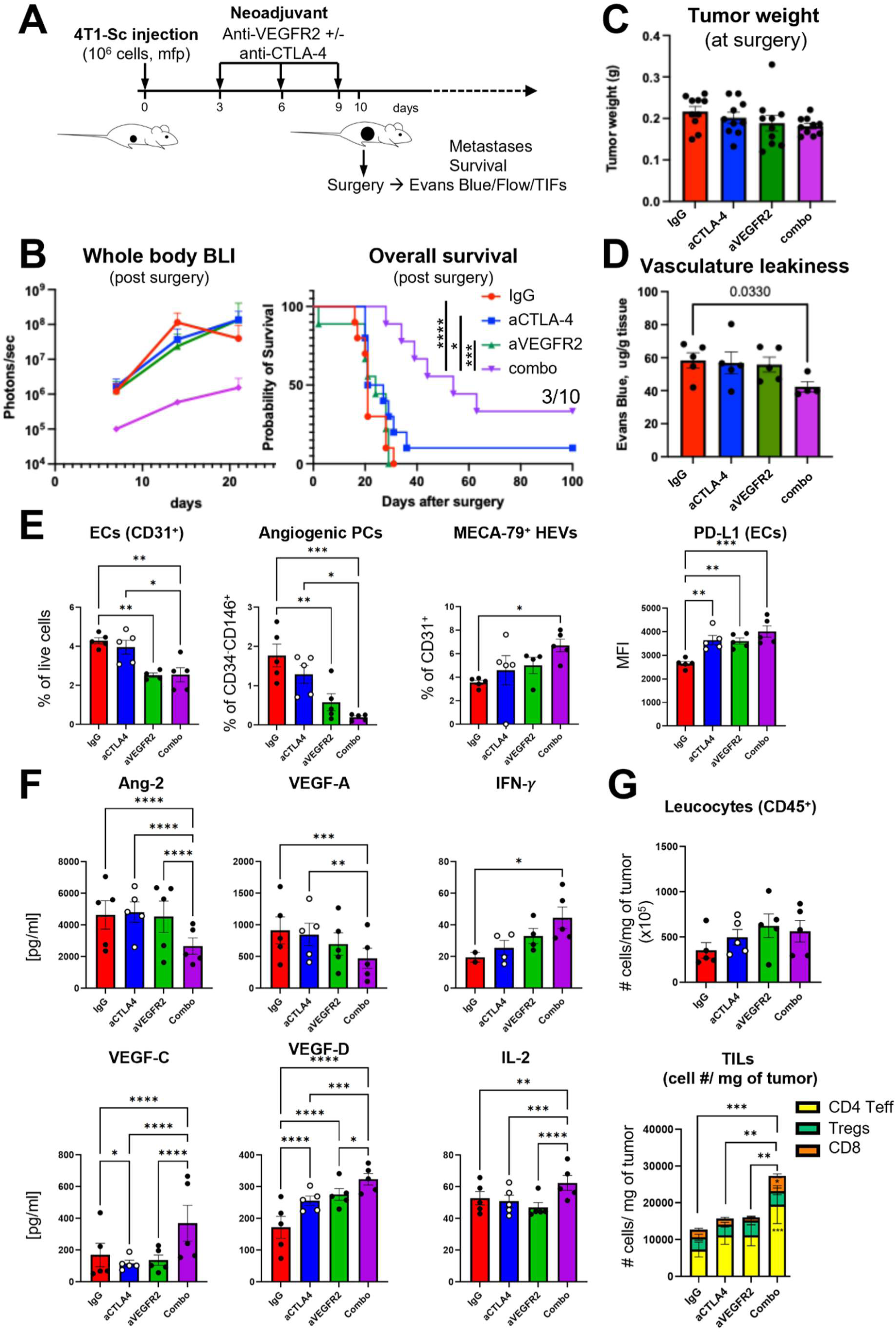
Anti-VEGFR2 restores the responsiveness of glycolytic tumors to anti-CTLA-4 immunotherapy. (**A**) Schema of neoadjuvant anti-VEGFR2 plus anti-CTLA-4 treatment in 4T1-Sc (glycolytic) tumor-bearing mice. (**B**) Metastasis progression after surgery by whole body BLI of Fluc (n=5/group) and overall survival (n=10/group from 2 independent experiments). (**C**) Tumor weight at the time of surgery after treatment as in (A). (**D**) Vasculature leakiness by Evans blue assay in mice treated as in (A). (**E**) Flow cytometry quantification of % CD31^+^ ECs, angiogenic PCs, HEVs and PD-L1 expression by MFI in ECs. (**F**) Quantification of the indicated soluble angiogenic and immune factors by Luminex beads-based immunoassay in TIF from tumors treated as in (A) and (**G**) flow cytometry quantification of intratumor CD45^+^ leucocyte and T-cell absolute numbers per mg of tumors in the same samples. Data are mean ± SEM of n=5-10 mice/group from 2 independent experiments. 2-sided unpaired *t* test: *, p<0.05; **, p<0.01; ***, p<0.001; ****, p<0.0001.

### Therapeutic, vascular, and immune effects following co-targeting of VEGFR2 and CTLA-4 depend on the tumor glycolytic state

We next asked if anti-VEGFR2 therapy in combination with anti-CTLA-4 had different activity depending on the glycolytic state of the tumor and underlying vasculature features. To this end, we compared the effects of this treatment against Sc and LDHA-KD tumors in the neoadjuvant setting using the 4T1 orthotopic model as in **Fig. 4A** and validated the results in the B16 orthotopic model.

In the 4T1 model, we observed that the combination treatment consistently improved metastases-free survival after surgery in mice that were implanted with 4T1-Sc; however, in the context of 4T1-KD tumors, addition of anti-VEGFR2 did not further delay metastatic dissemination compared to anti-CTLA-4 alone (**Fig. 5A**). In the primary tumor treatment setting with the B16 model, anti-CTLA-4 plus anti-VEGFR2 significantly reduced tumor growth and extended the survival of Sc-bearing mice but was completely ineffective against KD tumors (**Fig. 5B, C**). Moreover, the therapeutic effects of the combination therapy in B16-Sc-bearing mice were linked to reduced circulating tumor cells (CTCs) in peripheral blood, which were not significantly decreased by the same treatment in B16-KD tumor-bearing mice (**Fig. 5D**). This suggested the potential of anti-VEGFR2 with anti-CTLA-4 to improve the control of metastases according to the tumor glycolytic state in the B16 model as well.

**Figure 5:**
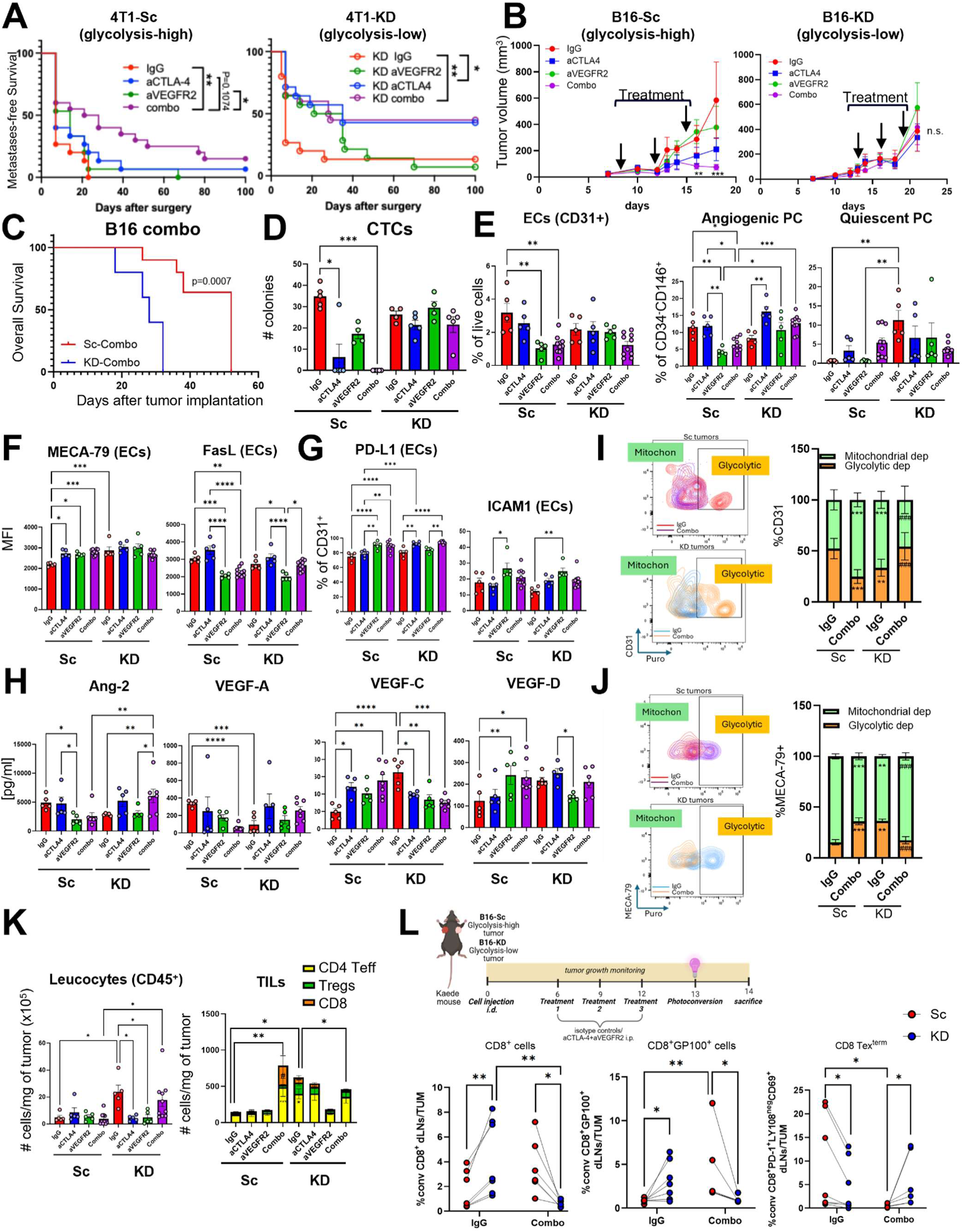
Improved activity of anti-VEGFR2 plus anti-CTLA-4 in glycolytic compared to glycolysis-defective tumors. (**A**) WT Balb/c mice were orthotopically injected in the m.f.p. with 4T1-Sc or 4T1-KD (1×10^6^ cells) and treated with neoadjuvant anti-CTLA-4 4 ± anti-VEGFR2 or isotype control as in Fig. 4A. Primary tumors were surgically resected on day 10 and 13 for 4T1-Sc and 4T1-KD respectively to ensure similar tumor burden was reached between the two models at the time of surgery. Mice were then followed for metastases by whole body BLI, and metastasis-free survival is reported for mice enrolled in 3 independent surgery experiments (pooled n=14-20/group). (**B-D**) WT C57BL/6 mice were orthotopically injected i.d. with B16-Sc or B16-KD (0.3×10^6^ cells) and treated with anti-CTLA-4 4 ± anti-VEGFR2 or isotype control as indicated by the arrows, and (B) primary tumor growth (n=5-10), (C) overall survival of combo-treated groups (n=10), and (D) circulating tumor cells at the end of treatment (n=5/each) were evaluated. (**E-K**) 4T1 tumors from mice treated as in (A) were processed for: flow cytometry quantification of frequencies of the indicated vascular cells (E), (F) expression (by MFI) of the indicated markers in ECs, and (G) frequencies of ECs expressing the indicated markers; TIF analyses by Luminex bead-based immunoassays to measure the concentration of the indicated hem/lymphangiogenic factors (H) (n=5-10/group); SCENITH analyses of ECs (I) and (J) HEVs (puromycin incorporation was used as a readout of ATP production under oligomycin treatment, where puromycin⁺ cells indicate glycolytic dependence and puromycin⁻ cells indicate mitochondrial dependence; n=8/group in 1 of 2 independent experiments); flow cytometry quantification of CD45^+^ leucocytes and tumor infiltrating lymphocytes (TILs: CD8^+^, CD4^+^Foxp3^-^Teff, CD4^+^Foxp3^+^Tregs) per mg of tumor (K; n=5-10/group, from 1 of 3 independent experiments). (**L**) Schema of treatment in Kaede mice implanted with contralateral B16-Sc and B16-KD tumors photoconverted to measure T-cell egress to dLN depending on the glycolytic state and treatment and flow cytometry quantification of ratio photoconverted CD8^+^ T cells, gp100-specific CD8^+^ T cells and terminally exhausted CD8^+^ T cells (Tex^term^; by surface staining of PD-1, CD69, and LY108 expression for compatibility with Kaede protein detection by surface staining) in dLN relative to tumor (TUM) in control vs. combo-treated mice (combined from 2 independent experiments with 2-3 and 4 mice/condition each, with similar results). Data are mean ± SEM. *, P<0.05; **, P<0.01; ***, P<0.001; ****, P<0.0001.

In the TME, as previously observed, KD tumors displayed improved vascular features vs. Sc tumors at baseline (control IgG treatment), including reduced numbers of ECs and angiogenic PCs and increased frequency of quiescent PCs, and similar profiles were restored in Sc tumors upon the combination treatment in both 4T1 and B16 models (**Fig. 5E** and **Suppl. Fig. 4A**). However, this beneficial vascular profile was not maintained in KD tumors upon the combination treatment, which appeared detrimental for vascular normalization in this tumor metabolic setting (**Fig. 5E** and **Suppl. Fig. 4A**). Further phenotyping of the vasculature revealed that, unlike in glycolysis-high tumors, the combination therapy did not modulate the presence of HEVs, nor did it decrease the frequency of FAS-L⁺, or consistently increased PD-L1⁺, ICAM-1⁺, or MHC-II⁺ ECs in glycolysis-low tumors (**Fig. 5F, G**; **Suppl. Fig. 4B**). Moreover, anti-CTLA-4 plus anti-VEGFR2 treatment decreased hemangiogenic factors (e.g., Ang-2, VEGF-A) and increased pro-lymphangiogenic factors VEGF-C and VEGF-D in Sc tumors, but not in KD tumors, where these factors were eventually modulated following the opposite trend (**Fig. 5H**, **Suppl. Fig. 4C**). These data indicate that the therapeutic combination opposed the vascular normalization effects driven by LDHA-KD in tumors, supporting the notion that this treatment is preferentially beneficial under conditions of dysregulated angiogenesis, as we observed in glycolysis-high tumors and not when the vascularization is already normalized, as in glycolysis-low tumors.

Importantly, the combination treatment profoundly modulated the metabolic state of vascular cells. CD31^+^ ECs from Sc tumors significantly decreased their glycolytic dependence and gained in mitochondrial dependence upon the combination therapy, while the opposite effects were observed with the same treatment in LDHA-KD tumors (**Fig. 5I**). Conversely MECA-79^+^ HEVs – which are more glycolytic in LDHA-KD vs Sc tumors at the steady state – acquired glycolytic dependence at the expenses of mitochondrial dependence in Sc tumors upon the combination treatment (**Fig. 5J**). Once again, the same treatment achieved the opposite effect in LDHA-KD tumors (**Fig. 5J**). It is possible that these vascular metabolic effects of the treatment occur as a consequence of the parallel changes in angiogenic factors, which follow opposite directions in Sc and LDHA-KD tumors (**Fig. 5H**).

We next asked if the vascular effects of the combination treatment also impacted T-cell recirculation in and out of the TME according to the tumor glycolytic state. The increased intratumor T-cell infiltration (especially with CD4^+^FOXP3^-^ T cells and CD8^+^ T cells) associated with vascular normalization in the combination-treated Sc tumors (**Fig. 4G**) was confirmed in both 4T1 and B16 glycolytic models (**Fig. 5K, Suppl. Fig. 4D**). However, in KD tumors, T-cell infiltration was decreased (**Fig. 5K)** or was not further augmented (**Suppl. Fig. 4D**) upon the combination treatment, potentially due to worsened vasculature and consequent impaired T-cell access to the tumor core in this condition. We next tested T-cell trafficking from the tumor to dLNs, which may serve to support systemic immunosurveillance against metastases. To this end, we employed Kaede C57BL/6 transgenic mice implanted with the syngeneic B16-Sc and KD-tumors in contralateral sides. Mice were treated with IgG or anti-CTLA-4 plus anti-VEGFR2 before tumor photoconversion and analysis of T-cell egress to dLN (**Fig. 5L**). According to our previous results with unilateral tumors (**Fig. 3N**), KD tumors showed greater CD8⁺ T-cell migration to dLNs than Sc tumors at the steady-state in IgG-treated mice (**Fig. 5L**). Moreover, tumor-egressing CD8⁺ T cells were enriched in tumor-antigen specific T cells recognizing the melanoma antigen gp100 (CD8⁺GP100⁺) and less likely displayed terminally exhausted phenotypes (PD-1⁺LY108⁻CD69⁺)^47^ in the context of KD vs. Sc tumors at the steady state (IgG control; **Fig. 5L**). Notably, the combination therapy completely reversed these effects: Sc-derived CD8⁺ T cells (including gp100-specific cells) from Sc tumors migrated more efficiently to dLNs, whereas KD tumors showed increased egress of exhausted CD8^+^T cells (**Fig. 5L**). Tumor size between KD and Sc tumors in these mice did not substantially differ, excluding this as a potential confounding factor (**Suppl. Fig. 4E**). Consistent with previous results with Kaede mice bearing unilateral KD or Sc tumors (**Fig. 3N**), we did not observe significant differences in tumor-dLN migration with respect to other immune cell subsets, including CD4⁺GITR⁻ T cells and CD4⁺GITR^hi^ Tregs, either with control or combination treatment (**Suppl. Fig. 4E** and not shown).

Overall, these results indicate that anti-VEGFR2 with anti-CTLA-4 promotes vasculature normalization linked to improved control of metastasis dissemination and CD8^+^ T-cell recirculation in glycolytic but not in glycolysis-defective tumors. These data point to vasculature abnormalities of glycolytic tumors as a therapeutic vulnerability and mechanism of resistance to immunotherapy, which can be overcome by normalizing the tumor vasculature, as we achieved here by using low-dose anti-VEGFR2 in combination with anti-CTLA4.

### Anti-VEGFR2 plus anti-CTLA-4 therapy mobilizes and activates central memory CD8^+^ T cells in glycolytic tumors

We next asked if T cells that recirculated intratumorally more abundantly in Sc tumors upon the combination therapy were also more activated and could directly mediate anti-tumor activity. We found that CD8^+^ T cells were indeed more activated (based on IFN-γ and TNF production) and less exhausted (based on Tox expression) in Sc tumors treated with anti-CTLA-4 plus anti-VEGFR2, while these parameters either did not change (B16 model; **Suppl. Fig. 4F, G**) or worsened (4T1 model; **Fig. 6A, B)** in glycolysis-defective tumors treated with the same regimen. Of note, the T-cell activation effect of the combination therapy in glycolytic tumors was restricted to CD8^+^ T cells, as CD4^+^Foxp3^-^ T cells did not increase (or eventually decreased) IFN-γ and/or TNF production capacity upon treatment (**Suppl. Fig. 5A**).

**Figure 6:**
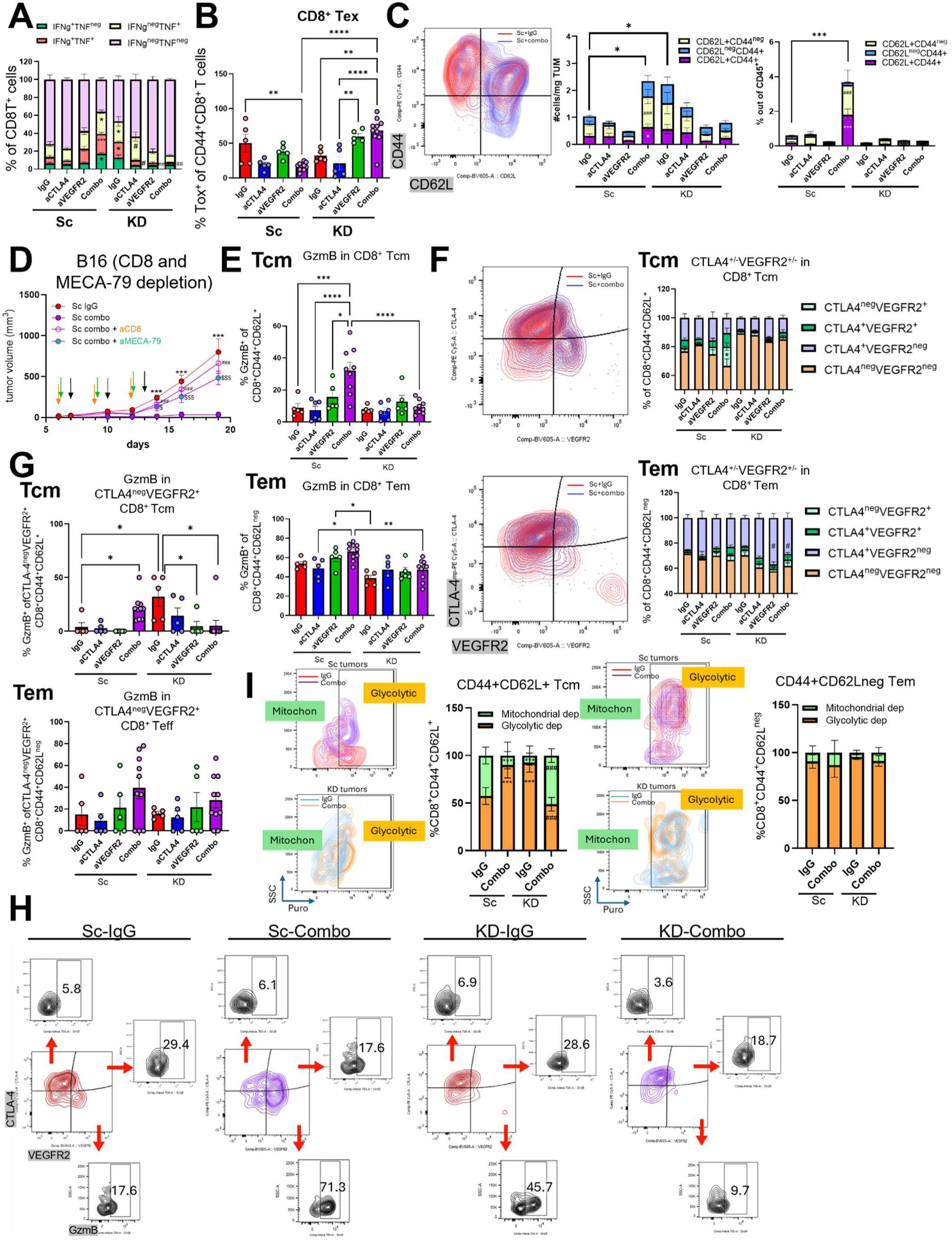
Intratumor T-cell functional changes upon anti-CTLA-4 plus anti-VEGFR2 treatment according to the tumor glycolytic state. 4T1-bearing mice were treated as in Fig. 5A and (**A**) frequencies of CD8^+^ T cells expressing IFN-γ ± TNF, and (**B**) frequencies of antigen-experienced CD44^+^CD8^+^ T cells expressing the exhaustion marker Tox were quantified by flow cytometry in tumors after treatment (n=5-10/group, from 1 of 3 independent experiments). (**C**) Representative flow cytometry plot of CD62L and CD44 in CD8^+^ TILs and quantification of CD8^+^ CD62L^+^CD44^-^ naïve, CD62L^+^CD44^+^, central memory (Tcm), and CD62L^-^CD44^+^ effector (Tem) cell frequencies and related absolute numbers per mg of tumors in 4T1-Sc and 4T1-KD tumors treated with anti-CTLA-4 ± anti-VEGFR2 or isotype control as in Fig. 5A. (**D**) Tumor growth profiles of glycolysis-high B16-Sc implanted in WT mice and treated with anti-CTLA-4 + anti-VEGFR2 (combo) alone or with a CD8 depleting (n=5/group) or a MECA-79 blocking antibody (n=5/group) in comparison with matched isotype control (n=5). Arrows indicate treatment time points. (**E-I**) Flow cytometry analyses of tumor-infiltrating CD8^+^ Tcm and Tem cells from 4T1-Sc and 4T1-KD tumors treated with anti-CTLA-4 ± anti-VEGFR2 or isotype control as in Fig. 5A: (E) quantification of granzyme B (GzmB)^+^ cell frequencies in CD8^+^ Tcm and Tem; (F) representative flow plots and quantification of CD8^+^ Tcm and Tem cell fractions expressing VEGFR2 ± CTLA-4; (G) quantification of GzmB^+^ cell frequencies within VEGFR2^+^CTLA-4^-^ CD8^+^ Tcm and Tem cell subsets and (I) representative flow plots of GzmB expression in VEGFR2 ± CTLA-4-expressing CD8^+^ Tcm subsets upon IgG control or combo treatment in 4T1-Sc and 4T1-KD tumors; (**I**) SCENITH analyses of CD8^+^ Tcm (left) and Tem (right) (puromycin incorporation was used as a readout of ATP production under oligomycin treatment, where puromycin⁺ cells indicate glycolytic dependence and puromycin⁻ cells indicate mitochondrial dependence). Data are mean ± SEM (n=5-10/group in one of 2-3 independent experiments). *, P<0.05; **, P<0.01; ***, P<0.001; ****, P<0.0001.

We thus investigated possible mechanisms that could drive T-cell infiltration and activation into Sc tumors with normalized vasculature upon anti-CTLA-4 plus anti-VEGFR2 vs. control treatment. In this condition in both tumor models tested (4T1-Sc and B16-Sc), the increased presence of HEVs was the most consistent effect (**Fig. 4E**, **5F, Suppl. Fig. 4B**). Given the role of HEVs in supporting T-cell recruitment in tissue through expression of peripheral node addressin (PNAd, detected by the antibody MECA-79) which binds CD62L^48^ on naïve and central memory T cells (Tcm), we examined infiltration of these T-cell subsets in comparison with CD62L^-^CD44^+^ effector memory T cells (Tem) in our tumor metabolic variants upon treatment. Sc tumors treated with the combination therapy showed a significant increase in CD62L⁺CD44^-^ naïve and CD44⁺CD8⁺ Tcm cells, compared to KD tumors in both the 4T1 and B16 models (**Fig. 6C; Suppl. Fig. 5B**). Overall, the quantitative and/or functional changes of CD8^+^ T cells and their CD62L^+^ fraction in glycolytic tumors treated with anti-VEGFR2 plus anti-CTLA-4 therapy were connected to the anti-tumor activity, as CD8^+^ T-cell depletion or MECA-79 blockade eliminated the therapeutic effect of this combination against glycolytic tumors (**Fig. 6D**).

Deeper phenotypic analysis to functionally characterize these CD62L^+^CD8^+^ T cells revealed that CD8⁺ Tcm cells infiltrating Sc tumors most strongly upregulate granzyme B (GzmB) upon the combination treatment (**Fig. 6E; Suppl. Fig. 5C left**). To clarify how this therapy enhanced the lytic activity of CD8^+^ Tcm cells, we asked if these cells could be directly targeted by anti-CTLA-4 and anti-VEGFR2 antibodies due to their expression of CTLA-4 and VEGFR2. Intriguingly, we observed selective expansion of CTLA-4⁻VEGFR2⁺ cells within CD8⁺ Tcm in Sc tumors upon either the combination treatment or anti-VEGFR2 alone (**Fig. 6F; Suppl. Fig. 5C, middle**) – a cell population that was not present among Tem cells or significantly increased in naïve cells after treatment (**Fig. 6F; Suppl. Fig. 5C, middle**). While the effects of CTLA-4 blockade on T cells have been widely investigated^49,50^, the effects of VEGFR2 blockade in T cells, especially CD8^+^ T cells, remain less clear. We thus compared VEGFR2 expression (MFI and frequency) across stromal cell subsets from our 4T1 and B16 tumor models. We found that VEGFR2 was expressed in the CD8^+^ T-cell compartment at levels comparable to vessel cells (**Suppl. Fig. 5D**), indicating the potential functional impact of VEGFR2 blockade in CD8^+^ T cells as well. Accordingly, tumor-infiltrating CTLA-4⁻VEGFR2⁺ CD8^+^ T cells, especially in the Tcm differentiation state, increased their cytotoxic phenotype (based on GzmB expression) upon the combination treatment in Sc tumors (**Fig. 6G-H; Suppl. Fig. 5C, right**). Consistently, CD8^+^ Tcm cells from combo-treated Sc tumors exhibited greater gain in *ex vivo* tumor-killing activity (**Suppl. Fig. 5E**) and glycolytic metabolism (**Fig. 6I**) compared to CD8^+^ Tem cells from the same tumors, suggesting that these CD8^+^ Tcm cells switch toward glucose metabolism to support rapid activation, proliferation, and effector function, as opposed to Tem cells that exhibit stable glycolytic dependence regardless of context, reflecting their more stable effector phenotype (**Fig. 6I; Suppl. Fig. 5E**). Of note, the combination of anti-CTLA-4 plus anti-VEGFR2 did not functionally modulate the same T-cell subsets in KD tumors (**Fig. 6E-H; Suppl. Fig. 5C**) and decreased the glycolytic activity of CD8^+^ Tcm in these tumors (**Fig. 6I**) according to their decreased *ex vivo* killing activity (**Suppl. Fig. 5E).** No significant or consistent differences were found in the maturation phenotype or activation state of tumor-infiltrating CD4⁺FOXP3⁻ T cells or Tregs across models and treatments (**Suppl. Fig. 5F** and not shown), further supporting a primary role for CD8^+^ T-cell trafficking and activation in the restoration of immune surveillance of glycolytic tumors upon vasculature normalization with anti-CTLA-4 and anti-VEGFR2 combination therapy.

### Anti-VEGF therapy in combination with ICB preferentially benefits patients with glycolysis-high tumors

Inhibition of the VEGF pathway with the anti-VEGF bevacizumab in combination with the PD-L1 ICB agent atezolizumab is approved in patients with hepatocellular carcinoma (HCC) without any additional standard therapy. There are no other ICB therapies currently approved with anti-VEGF or anti-VEGFR2. We thus leveraged a public tumor transcriptomic data set from HCC patients treated with atezolizumab alone in comparison with atezolizumab with bevacizumab (GO30140 group F)^38^ to test in patients the relevance of targeting angiogenesis with ICB depending on the tumor glycolytic state. To this end, we stratified patients in glycolysis-high and glycolysis-low groups based on the median glycolysis signature score of their tumors (**Suppl. Table 1**) and then compared overall survival between bevacizumab plus atezolizumab and atezolizumab alone in patients within each group. We found that patients with glycolysis-high tumors experienced significantly improved survival if they were treated with the combination therapy compared to ICB alone (**Fig. 7**). In contrast, patients with glycolysis-low tumors did not gain any substantial benefit from the combination treatment with respect to ICB alone (**Fig. 7**). These results in the human setting reflect our preclinical data comparing glycolysis-high (Sc) and glycolysis-low (KD) tumor mouse models and support the relevance of measuring the tumor glycolytic state to identify patients that can most greatly benefit from anti-angiogenic therapy in combination with immunotherapy.

**Figure 7:**
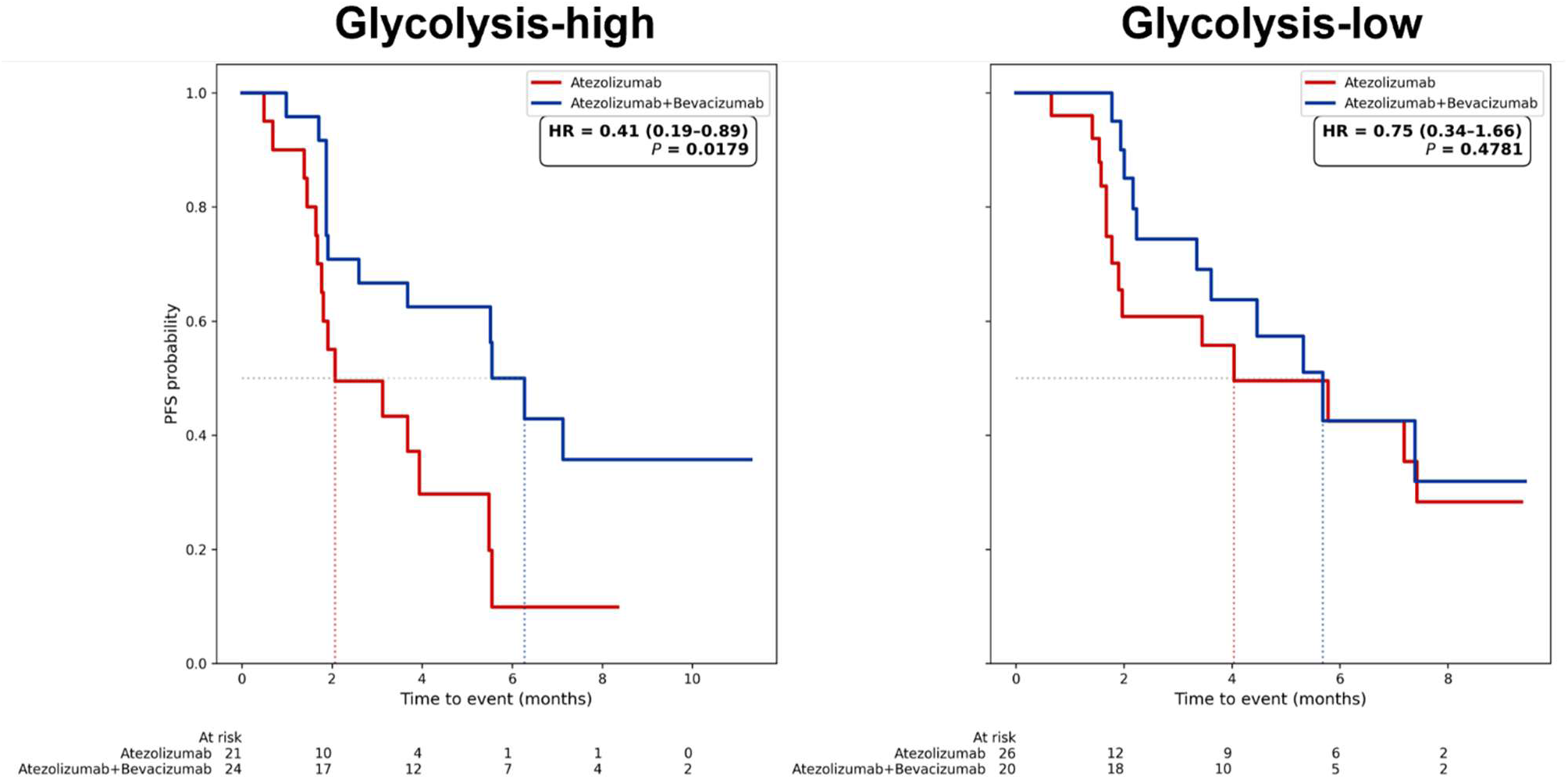
Antiangiogenic therapy enhances the efficacy of immunotherapy selectively in glycolysis-high tumors. HCC patients in group F of the GO30140 Phase Ib trial testing atezolizumab alone vs. atezolizumab plus bevacizumab^38^ were stratified in glycolysis-high and glycolysis-low based on the median glycolysis signature (Suppl. Table 1) score of their tumor transcriptomes (EGAS00001005503), and survival (PFS) was compared between the two treatments within each glycolysis group. HR p values are reported on each plot.

## Discussion

The tumor vasculature is emerging as a critical component regulating local and systemic anti-tumor T-cell responses, making it crucial to determine how to modulate tumor angiogenesis for improved immunotherapy^51^. Our work addresses this by mechanistically connecting abnormal angiogenesis and T-cell exclusion with the glycolytic state of tumor cells. This provides a novel biological framework for rationally combining immunotherapy and vasculature normalization approaches based on the tumor metabolic state.

We found that lowering glycolysis in cancer cells improves the vasculature in the TME in terms of phenotypic, metabolic, and functional parameters, by (1) reducing ECs and progenitor ECs, (2) rewiring the phenotype of PCs toward quiescent stabilizing PCs, (3) inducing the upregulation of immune interacting molecules and HEV phenotype while downregulating T-cell pro-apoptotic molecules in ECs, (4) modulating metabolic dependencies in ECs and HEVs (reducing glycolytic metabolism in ECs while increasing it in HEVs), (5) increasing vessel PC coverage, (6) reducing vasculature leakiness and HIF-1α expression/activity in the TME. Importantly, these tumor metabolism-driven vasculature effects made the TME permissive to T-cell trafficking, function, and survival. Notably, we found that correcting the tumor vasculature in glycolytic tumors using low-dose anti-VEGFR2 restored the responsiveness of these otherwise immune-refractory tumors to CTLA-4 blockade by restoring vasculature and immune features of glycolysis-low tumors. Mechanistically, the anti-VEGFR2 plus anti-CTLA-4 combination increased intratumoral MECA-79^+^ HEVs and associated recruitment of CD62L^+^CD8^+^ T cells in glycolytic tumors, with CD8^+^ Tcm cells becoming activated and maturing into cytotoxic GzmB^+^VEGFR2^+^CTLA-4^-^CD62L^+^CD44^+^CD8^+^ Tcm cells potentially through their glycolytic switch. Intriguingly, the same combination therapy opposed the vascular and immune benefits of LDHA-KD glycolysis-defective tumors observed at the steady state.

These results uncover several novel concepts. First, our findings unravel the mechanistic implications of tumor cell glycolysis in the TME and response to immunotherapy through vasculature remodeling. Prior data, including from our group, had established the effect of lactate – the end-product of glycolysis – to induce VEGFA and ANGPT2 expression by activating HIF-1 and the hypoxia response program in cancer cells^22,23^. Separately, the impact of hypoxia in driving immune dysfunction in the TME has been extensively investigated^52^. Here, we demonstrate that these mechanisms are interconnected and regulated at the level of tumor cell glycolysis. We find that dampening glycolysis in cancer cells by LDHA-KD normalizes the TME at the vascular and immune levels and that similar effects can be achieved by pharmacologic blockade of both VEGFR2 and CTLA-4 in glycolytic tumors.

Second, our data suggest that vasculature rewiring based on the tumor glycolytic state involves both hemangiogenesis and lymphangiogensis. We find that in the TME formed by LDHA-KD vs. glycolytic cancer cells, the concentration of lymphangiogenic factors VEGF-C and VEGF-D increases, whereas the concentration of hemangiogenic factors VEGF-A and ANG-2 decreases. While the precise source of these factors in the TME remains unclear, LDHA-KD glycolysis-low tumor cells decrease their production of VEGF-A ^22^ (**Suppl. Fig. 2D**). These findings suggest that the glycolytic state of cancer cells may directly shape their angiogenic potential, thereby influencing endothelial metabolism and the resultant vascular phenotype. Intriguingly, similar rewiring of hem/lymphangiogenic factors in the TME is observed in glycolytic tumors upon anti-VEGFR2 plus anti-CTLA-4 therapy, which do not directly modulate cancer cells. Changes in these factors for angiogenesis signaling in the TME may shape the tumor vasculature by impacting vessel cell metabolism. Accordingly, ECs derived from LDHA-KD tumors or from glycolytic tumors treated with anti-VEGFR2 plus anti-CTLA-4 exhibited significantly reduced glycolytic dependence, reflecting a metabolic shift toward a more oxidative, quiescent, and stabilized EC phenotype that could support vascular normalization and permissive immune infiltration^53,54^. In striking contrast, tumor-associated HEVs induced under these same metabolic and treatment conditions showed enhanced glycolytic activity – a metabolic program that could sustain their expansion. Moreover, LDHA-KD in cancer cells or the combination treatment in glycolytic tumors increased the egress of CD8^+^ T cells (including tumor-antigen-specific T cells) from the TME to the dLN, which was associated with greater protection from metastasis progression. This result suggests that the vasculature normalization we observe in the same treatment conditions may extend to both blood and lymphatic vessels and that their cooperation is necessary to restore full T-cell recirculation in and out of the tumor for local tumor destruction and systemic immunosurveillance, respectively. Specifically, tumor-infiltrating T cells recirculating to the dLN might improve surveillance of metastases, as T cells that egress tissue exhibit effector function^55^ and can seed protective LN resident memory^56,57^. Similar to the blood vasculature, recent work demonstrates that the lymphatic vasculature can undergo a normalization process that limits metastasis and improves response to ICB^58^. Lymphatic normalization can be driven by local cytotoxic T-cell activity and depends upon metabolic reprogramming, notably the suppression of oxidative phosphorylation^58^. In the future, it will be important to elucidate the precise mechanisms through which tumor metabolic rewiring alters tumor lymph/hemangiogenic potential and re-shapes blood and lymphatic networks. Tumor-derived metabolites can impact stromal cells in the TME and also dLNs. Relevant to our findings, tumor-derived lactate was found to trigger the angiogenic phenotype of intratumoral ECs^59^ and to also contribute to the dLN metastatic niche^60^. These mechanisms could play a role in mediating the different vasculature phenotypes and metastatic potential of LDHA-KD vs. glycolytic tumors reported here.

Third, our study provides novel opportunities to guide improved use of anti-angiogenic therapy for tumor vasculature normalization in combination with immunotherapy. The concept that anti-angiogenics should be dosed with the intent of normalizing the vasculature rather than destroying it, as a means to enhance drug delivery when used in combination regimens, was proposed more than 20 years ago^61^. This concept becomes extremely important in the context of immunotherapy, which relies on immune cell trafficking into the TME. In addition, given the inherent risk of tumor vasculature normalization to potentially facilitate tumor cell growth and/or dissemination, coupling this intervention with means to enhance immunosurveillance appears to be a logical approach to pursue further. However, how to precisely achieve tumor vasculature normalization for improving intratumor T-cell trafficking and function is a major challenge. Our findings indicate that the tumor glycolytic state is a key determinant of tumor vasculature normalization. In mouse models, we found that tumor cell glycolysis influences tumor angiogenesis and intratumor T-cell trafficking. In addition, our results suggest that tumor vasculature normalization supports the therapeutic activity of immunotherapy only in the context of highly glycolytic tumors (**Suppl. Fig. 6A**). In human tumors, we found that tumor glycolysis positively correlates with abnormal vasculature and with poor immune cytolytic activity, and that the combination of these features predicts outcomes across most aggressive and immunotherapy-resistant tumor types. Overall, this supports the rationale to dose anti-angiogenics based on the tumor glycolytic state. To implement such an approach in the clinic, it will be critical to determine practical means to infer the tumor glycolytic state. Baseline non-invasive metabolic imaging using [^18^F]FDG/PET, coupled with longitudinal DCE MRI, could be tested in patients to assess the relationship between these metabolic and vasculature parameters and in association with response to current regimens of anti-angiogenics in combination with immunotherapy. This could then support the use of the same non-invasive imaging approaches to guide and refine these treatments prospectively in patients in the future.

Fourth, we report previously unrecognized synergistic effects of VEGFR2 and CTLA-4 blockade on both the tumor vasculature and T cells. Specifically, we found that anti-VEGFR2 drives most of the vasculature-normalizing effects when administered together with anti-CTLA-4, indicating the cooperation of these independent axes on the tumor vasculature. Prior studies had shown that intratumor effector T cells play a critical role in vasculature normalization, which can be promoted by ICB therapy^42,62–64^. Here, we observed that the combination of anti-VEGFR2 and anti-CTLA-4 increases MECA-79^+^ HEVs in glycolytic tumors, triggering infiltration by CD62L^+^CD8^+^ T cells. A recent report described the effect of CTLA-4 blockade to enrich HEVs in the tumor by depleting tumor-associated ECs^42^. In our glycolytic tumor models, we observed similar decreases of ECs upon anti-VEGFR2 with or without anti-CTLA-4, and selective HEV increases upon the combination therapy, suggesting CTLA-4 blockade could contribute additional mechanisms driving the HEV phenotype. For example, CD4^+^ T cells have been implicated in the induction of HEVs in tumor^42^. Specifically, independent studies reported the effect of Treg depletion to promote HEV formation^65–67^. In our work, we utilized a CTLA-4 blocking antibody that does not substantially decrease intratumoral Tregs, as previously reported and also documented by our data (**Fig. 4G**; **Fig. 5K**; **Suppl. Fig. 4D**), suggesting that alternative mechanisms may be involved. Notably, in human tumors, the presence of HEVs at baseline has been found to be associated with better outcomes, particularly in patients treated with anti-CTLA-4-based immunotherapy^42^. HEVs are often found associated with tertiary lymphoid structures (TLSs)^37,68^, which represent ectopic lymphoid aggregates, including lymphocytes, dendritic cells (DCs), and other innate immune cells. The presence of TLSs in human tumors has also been reported to correlate with a positive outcome to immunotherapy^69–71^. In these structures, HEVs represent critical entry sites for immune cells, especially anti-tumor T cells. Our data show that tumor glycolysis negatively correlates with the presence of HEVs and that integrating HEV features within a signature incorporating key tumor metabolic, vasculature, and immune components of the TME better predicts PFI in most tumor types in the TCGA. These results illustrate the potential role of tumor glycolysis in restraining TLS formation by promoting vascular abnormalities, supporting further research to elucidate the mechanistic link between these processes.

As a novel on-T-cell synergistic effect of anti-CTLA-4 plus anti-VEGFR2, we observed expansion and activation of VEGFR2^+^CTLA-4^-^CD8^+^ Tcm cells upon the combination treatment in the TME of glycolytic tumors responding to therapy. This suggests that enhanced CD8^+^ Tcm cell recruitment – potentially driven by HEV formation – is paired with functional modulation of the same T cells in the TME by anti-CTLA-4 and anti-VEGFR2 (**Suppl. Fig. 6B**). Intriguingly, these CD8^+^ Tcm cells acquired a glycolytic phenotype in the TME of glycolytic tumors upon anti-CTLA-4 plus anti-VEGFR2 treatment – effect that could potentially sustain their anti-tumor function. Previous work has shown that VEGFR2 is expressed in highly suppressive Foxp3^hi^ Tregs and signals for Treg proliferation^72,73^, and that VEGF-A:VEGFR2 blockade can limit Treg expansion, reinvigorates CD8⁺ T cells, and reduces exhaustion^74,75^. In addition, VEGFR2 expression was reported to promote upregulation of immune checkpoint molecules in CD8^+^ T cells^75,76^ linked to TCR signaling inhibition^77^. A more recent study reported an increase of VEGFR2^+^ CD4^+^ Th1 and CD8^+^ T cells in peripheral blood of colon cancer patients compared to healthy controls^78^. These previous findings suggest that the VEGF:VEGFR2 pathway may contribute to inhibiting anti-tumor CD8^+^ T cells. Accordingly, a recent report showed that anti-VEGFR2 treatment can enhance CD8⁺ T-cell functionality and migratory capacity^79^. This supports our observations that VEGFR2 blockade, in combination with CTLA-4 blockade, expands and functionally reinvigorates VEGFR2^+^CD8^+^T cells within the TME. Importantly, we observed these effects in highly glycolytic tumors, but not in glycolysis-defective LDHA-KD tumors, where VEGFA is less abundant (**Fig. 5H; Suppl. Fig. 4C**) and the VEGFR2 pathway may thus be less active. This suggests once again the importance of monitoring the tumor glycolytic state and related angiogenic pathways to appropriately intervene therapeutically. Proper modulation of tumor vasculature based on the tumor glycolytic state to normalize the TME may be particularly relevant to enhance intratumor recruitment and efficacy of T-cell therapies, which are becoming a central component of the immunotherapy arsenal. In support of such an approach, initial evidence shows the negative impact of tumor glycolysis in the response to TIL therapy in patients^80^. Moreover, given our observations that tumor vasculature normalization also restores tumor-specific T-cell trafficking from the TME to the periphery, approaches to vascular normalization may be exploited in the context of neoadjuvant therapies to promote systemic T-cell-mediated tumor immunosurveillance after surgical resection of primary tumors.

Taken together, our work contributes to resolving the complexity of TME physiology by demonstrating the mechanistic role of tumor metabolism in shaping vasculature and T-cell response dynamics. Our data identify aberrant angiogenesis as a rational vulnerability of glycolytic tumors that can be targeted to enhance anti-tumor T-cell response. In addition, these results demonstrate the importance of normalizing the TME physiology to restore T-cell trafficking and activation and enable long-lasting anti-tumor immunity.

## Materials and Methods

### Tumor cell lines

The mouse mammary carcinoma 4T1 cell line and the mouse melanoma B16F10 cell line (B16) were provided, respectively, by F. Miller and I. Fidler. Tumor cells were transfected with SureSilencing™ LDHA-targeting shRNA plasmids (KD; A2= GTACGTCCATGATGCATATCT;) or scramble control plasmids (Sc= GGAATCTCATTCGATGCATAC) (QIAGEN). Stable LDHA-KD (4T1 A2–10KD = 4T1-KD, and B16-KD) and scramble control (4T1-Sc and B16-Sc) cell lines were generated as previously described^13,22^. The 4T1 cell lines are also transduced with Firefly luciferase (Fluc) and the exGaussia luciferase (exGluc) HRE-reporter vectors, as previously described^22^. 4T1 cell lines were cultured in DMEM supplemented with 10% heat inactivated FBS, 25 mM glucose, 6 mM L-glutamine, 1 x penicillin/streptomycin, 500 mg/L gentamycin (G418), and 4 mg/L puromycin. The addition of these antibiotics is required to support the growth of selected cells bearing the HRE-exGluc reporter and shRNA bearing cassette. B16 cell lines were cultured in RPMI1640 supplemented with 10% inactivated FBS, 1× nonessential amino acids, and 2 mM l-glutamine, and selected with 4 mg/L puromycin.

LDHA downregulation in LDHA-KD vs. Sc cell variants was routinely tested and confirmed at the protein level by western blotting, using a rabbit anti-LDHA antibody (1:1,000; Cell Signaling Technology, cat# 2012S) coupled with an HRP-conjugated anti-rabbit IgG (1:2,000; Cell Signaling Technology, cat#7074S) as secondary antibody, along with beta actin (1:5,000, Sigma, cat#A2103) as protein loading control, and at enzymatic activity level by using the Cytotoxicity Detection Kit PLUS (LDH) (Roche Diagnostics), as previously reported^13,22^.

### Patient samples

Primary human TNBC FFPE samples were obtained from the Weill Cornell Medicine tissue archive under IRB #1603017108. All patients signed an approved informed consent before any study-related procedures.

### Seahorse analyses

Differences in glycolytic capacity between LDHA-KD and Sc tumor cell lines were measured by Glycolytic Proton Efflux Rate and ATP rate assays using a Seahorse XF 96 Analyzer according to the manufacturer’s instructions (Seahorse XF Glycolytic Rate Assay and Real-Time ATP Rate Assay, Agilent Technologies).

For Seahorse analyses of tumor-derived ECs, tumors were mechanically dissociated and digested enzymatically, and the resulting suspension was passed through a 70 µm strainer. Viable cells were enriched by a 40% Percoll gradient, and the EC fraction was subsequently isolated using CD31 MicroBeads (Miltenyi) according to the manufacturer’s protocol. Purified ECs were seeded on XF96 Seahorse plates (Agilent) pre-coated with collagen (10 µg/mL, O.N., 4 °C) at 2×10^4^ cells/well and allowed to adhere overnight in endothelial growth medium (PromoCell). mrVEGF-A (PeproTech) was added at 10 ng/ml and cultures were incubated for ∼18hrs. On the assay day, medium was replaced with Seahorse XF assay medium (non-buffered DMEM) supplemented with 10 mM glucose, 1 mM pyruvate, and 2 mM glutamine (pH 7.4), and plates were incubated 45–60 min in a non-CO₂ incubator. OCR and ECAR were measured using Glycolytic Stress Test. Data were analyzed using Wave (Agilent) and normalized to cell number or protein content as indicated in the figure legends.

### In vivo experiments

4T1 syngeneic female BALB/cAnN mice were from Charles River Laboratory, RAG2 KO BALB/c from Taconic, and B16 syngeneic WT C57BL/6 mice from Jackson Laboratory. Kaede C57BL/6 transgenic mice were available through Dr. Amanda Lund’s lab (New York University, NY, NY). All mice were bred and maintained under specific pathogen-free conditions (with a 12 h light-dark cycle at a temperature of 21–23°C and humidity of 35–55%) and used at the ages of 5–10 weeks.

For neoadjuvant treatment with the 4T1 models, 1 × 10^6^ tumor cells were injected orthotopically in the m.f.p. of 5-6-week-old female mice. Three days later, the tumor burden was quantified by BLI, and mice were randomized into different treatment groups. 4T1-Sc and 4T1-KD tumors were resected on day 10 and day 13 (3 days apart) after tumor implantation to equalize tumor sizes before surgery at a time point of tumor growth that precedes detection of metastases. Metastasis development was monitored weekly by BLI, as described below. For primary tumor treatments with the B16 models, 0.3 × 10^6^ tumor cells were implanted orthotopically intradermally (i.d.) in 5–8-week-old female mice, and treatment was started when tumors became palpable (day 7, B16-Sc; day 10, B16-KD). Primary tumor growth was measured twice a week by caliper.

To establish mice with similar-sized bilateral tumors at the time of adoptive cell transfer, 4T1-KD and 4T1-Sc cells (0.25×10^6^) were injected orthotopically in RAG2 KO mice 3 days apart.

Antibody treatments were performed by intraperitoneal injection of 100 μg anti-CTLA-4 (clone 9D9 IgG2b, BioXcell), 200 µg anti-VEGFR2 (clone DC101, BioXcell), their combination, or isotype controls (clone MPC-11 and clone HRPN, respectively, BioXcell) 3 days apart for three administrations as indicated in the figures. For CD8 depletion and MECA-79 blockade experiments, anti-CD8 (clone 2.43, Bioxcell) and anti-MECA-79 (clone MECA-79, Biolegend) were injected intraperitoneally (250 µg/mouse, anti-CD8) and intratumorally (15 µg/mouse, anti-MECA-79) respectively, one day before each treatment administration as indicated in figure.

For adoptive cell transfer in the bilateral 4T1 tumor model system, anti-tumor T cells were isolated from the spleens and lymph nodes of BALB/cAnN mice that had been serially immunized with the corresponding tumor model. T cells were activated with plate bound anti-CD3 (1 µg/ml) and anti-CD28 (5 µg/ml) for 72 hours and further expanded in IL-2 (10 U/ml), IL-7 (2.5 ng/ml), and IL-15 (25 ng/ml) for additional 24 hours before adoptive transfer into tumor-bearing RAG2 KO mice. T cells from 4T1-KD and 4T1-Sc immunized mice were mixed at a 1:1 ratio before infusion (1 × 10^6^ cells, i.v.).

To track immune cells egressing from primary tumors, Kaede transgenic mice were injected i.d. with 0.3 × 10⁶ B16-Sc and B16-KD in contralateral flanks. Tumor growth was monitored daily, and tumors were photoconverted after 12 days. Antibody treatments (as described above) were performed starting on day 6, 3 days apart, 3 times, before photoconversion. For photoconversion, tumors were exposed to violet light (wavelength 350–410 nm) using a cold-light source for 5 minutes under anesthesia, as previously described^46^. Twenty-four hours after photoconversion, mice were sacrificed, and tumors, dLNs, and spleens (as a control) were harvested for flow cytometry analysis to track Kaede-red⁺ (photoconverted) cells across tissues.

In all experiments, tumor growth was monitored every 2–3 days by caliper, and tumor volume was calculated using the formula: (length × width × depth)/2 (in cases where depth is measurable) or (longest dimension × shortest dimension × short dimension)/2.

The maximal tumor size of 20 mm in any direction was not exceeded in any experiment. All animal experiments were conducted according to protocols approved by the Weill Cornell Medicine and New York University Institutional Animal Care and Use Committee (IACUC) and carried out in accordance with institutional guidelines.

### Flow cytometry analyses

Tumors were dissociated after 30-minutes incubation at 37°C with Liberase TL and DNAse I (Roche) to obtain single-cell suspensions. When the tumor mass exceeded 0.1 g, immune-cell infiltrates were enriched using Percoll (GE Healthcare) gradient centrifugation. Surface staining was performed after 10 min incubation on ice with an anti-mouse CD16/CD32 antibody (clone 2.4G2, BD Biosciences) to block Fcγ receptors, by using panels of appropriately diluted fluorochrome-conjugated antibodies (from BD Biosciences, eBioscience, Invitrogen, or Biolegend; **Suppl. Table 2**). For intracellular staining, mouse cells were fixed and permeabilized using Foxp3 fixation/permeabilization buffer (eBioscience) according to the manufacturer’s instructions and then incubated with appropriately diluted antibodies (**Suppl. Table 2**).

For intracellular cytokine staining, cells were stimulated using the Cell Activation Cocktail (with Brefeldin A; BioLegend, Cat# 423303) according to the manufacturer’s instructions. Briefly, cells were resuspended in complete RPMI-1640 medium supplemented with 10% FBS and plated at a density of 1–2 × 10⁶ cells per well in a 96-well round-bottom plate. The activation cocktail was added directly to the culture at a final concentration of 1×, and cells were incubated for 4 hours at 37°C in a humidified incubator with 5% CO₂. Following stimulation, cells were washed with PBS and processed for surface and intracellular marker staining as described above.

To identify gp100-specific CD8⁺ T cells, staining with H-2Db/gp100_25–33 tetramer (KVPRNQDWL; MHC class I, APC-conjugated; obtained from the NIH Tetramer Core Facility) was performed. Briefly, single-cell suspensions were incubated with tetramer diluted at 1:500 in FACS buffer (PBS, 2% FBS, 2 mM EDTA) for 30 minutes at room temperature in the dark. After tetramer staining, cells were subjected to surface staining as described above.

For SCENITH^44^, freshly isolated cells were incubated for 30 min at 37 °C with puromycin (10 µg/mL) in the presence or absence of metabolic inhibitors: 2-deoxy-D-glucose (2-DG, 50 mM), oligomycin (1 µM), or their combination. Cells were subsequently stained with viability dye and surface markers, fixed, permeabilized, and intracellularly stained with anti-puromycin antibody to quantify protein synthesis as a proxy of ATP production. Samples were acquired by flow cytometry, and metabolic parameters (glucose dependence, mitochondrial dependence, glycolytic capacity) were calculated according to the SCENITH workflow^44^.

Cells were acquired using the BD FACSymphony A3 with BD FACSDiva and data analyzed using the FlowJo (10.10.0) software.

### FACS-sorting of TIL subsets and killing assay

IgG- and combo-treated Sc and KD tumors were excised, minced, and enzymatically digested as previously described. TILs were enriched by density gradient centrifugation (40/80% Percoll) and stained with fluorochrome-conjugated antibodies (CD45, CD4, CD8, CD62L, and CD44). Dead cells were excluded using DAPI. Following compensation and appropriate gating strategies, CD8⁺ TIL subsets (Tem: CD62L⁻CD44⁺ and Tcm: CD62L⁺CD44⁺) were sorted using a FACSAria II (BD Biosciences) under sterile conditions at >98% purity. Sorted populations were immediately processed for killing assays.

Sorted TILs were co-cultured with the corresponding target tumor cells at the indicated effector-to-target (E:T) ratio (typically 10:1) in complete T cell medium (RPMI-1640 supplemented with 10% FBS, L-glutamine, penicillin/streptomycin, sodium pyruvate and 50 µM beta-mercaptoethanol). Tumor cells were pre-treated for 18 hrs with IFN-γ (100 ng/mL) to increase MHC expression. Co-cultures were incubated for 24 hrs at 37°C in a humidified incubator. At the end of the assay, T cells were removed by gentle washing and the remaining tumor cells were trypsinized, counted, and replated at 1:100 dilution in 6-well plates in complete DMEM medium for colony formation. Cells were cultured for 7 days to allow surviving tumor cells to form colonies. Colonies were then fixed with cold methanol (10–15 min), stained with 0.1% crystal violet, rinsed, and air-dried. Colonies were counted manually. Tumor cell killing was expressed as the % reduction in colony number in wells exposed to T cells relative to tumor cells cultured without T cells, and reported as ratio of % killed tumor cells with T cells from the combo-treated group relative to the IgG-treated group.

### Immunofluorescence staining and quantification

Tumor tissue samples were fixed in buffered formalin (4% paraformaldehyde in 10 mM phosphate buffer, pH 7.4) for 24 hours at 4°C. Samples were then extensively washed and stored in histology cassettes at 4°C in 70% ETOH in H_2_O for 24-48hrs before embedding. Paraffin-embedded tissues were sectioned at 5 μm and baked at 58°C for 1 hr. Multiplex immunofluorescent (IF) staining was performed at the Molecular Cytology Core Facility of Memorial Sloan Kettering Cancer Center using the Leica Bond RX staining system. Slides were loaded in Leica Bond, and IF staining was performed as follows. Samples were dewaxed and pretreated with an EDTA-based epitope retrieval ER2 solution (Leica, cat# AR9640) for 20 minutes. at 100°C. Antibody staining and detection were conducted sequentially. Primary antibodies against HIF-1α (0.5 µg/ml, Novus Bio, cat#NB100-479), NG2 (5 µg/ml, Millipore, cat#AB5320), and CD31 (0.08 µg/ml, Abcam, cat#ab182981) were incubated for 1 hour, followed by 8 minutes incubation with Leica Bond Polymer anti-rabbit HRP (included in the Polymer Refine Detection Kit (Leica, cat#DS9800). Rabbit anti-mouse linker (Leica Bond Post-Primary reagent included in Polymer Refine Detection Kit, cat# DS9800) was incubated for 8 minutes before the application of the Leica Bond Polymer anti-rabbit HRP. After that, CF® dye tyramide conjugates 430, 594, and 750 (Biotium, Cat#96053, 92174, 96052) were used for detection. After each round of IF staining, epitope retrieval was performed for denaturation of primary and secondary antibodies before another primary antibody was applied. After the run was finished, slides were washed in PBS and incubated in 5 μg/ml 4’,6-diamidino-2-phenylindole (DAPI) (Sigma Aldrich) in PBS for 5 minutes, rinsed in PBS, and mounted in Mowiol 4–88 (Calbiochem). Slides were kept overnight at −20°C before imaging.

All segmentation and quantification analyses were performed in QuPath^81^ and exported in “csv” format for subsequent analysis in MATLAB and/or Prism. Briefly, images were color deconvolved with standard unmixing matrices and aligned with IF data as previously described^82^. For cell segmentation, DAPI extracted nuclear masks were expanded to 3 µm to generate cell masks. Positive cells and cell-colocalization were computed with custom MATLAB scripts based on exported QuPath measurements (for the nucleus alone or the entire segmented cell, depending on marker localization) using manually determined thresholds per channel. Multiple positive cell classes were computed as the AND of individual single positive markers on the same cell. Clustering analysis was implemented to identify spatial clusters in the union of NG2^+^ or CD31^+^ cells representing vessel structures. For each image, all positions for cells positive for either marker are extracted, and single linkage clustering is performed on x,y position using Euclidean distance with cutoff 20 µm. Each cluster, representing a vessel, was classified based on the relative composition of CD31^+^ and NG2^+^ cells. A cluster is considered positive for a marker if at least 10% of cells are positive for that marker (which was determined based on the distribution of NG2^+^ cells colocalizing with CD31^+^ cells in normal tissue). This defines 3 cluster classes (CD31^+^, NG2^+^, [CD31^+^NG2^+^]), though a priori, all vessels are expected to be CD31^+^. Clusters in each class are tallied, excluding singleton clusters. As a metric for vessel complexity, the mean count of CD31^+^ cells in vessel clusters from each image was computed.

### TIF analysis

For 4T1, freshly excised tumor samples were placed in a 1.5 ml tube and then centrifuged at 10,000 × g for 15 minutes at 4°C. For B16, tumors were placed on a 25 ml centrifuge filter and centrifuged at 8,000 × g for 15 minutes. The fluid was collected at the bottom of the tube and stored at −80 °C for analysis. TIFs were diluted at 1:10 with Multiplex Assay Buffer (Millipore) for quantification of soluble protein factors by Luminex-based bead multiplex immunoassays (Millipore) using the DropArray system (Curiox BioSystems), according to the manufacturer’s instructions.

### Evans Blue assays

Vascular permeability was assessed using Evans blue assays. Evans blue, which binds to serum albumin, was administered to tumor-bearing mice by retro-orbital injection (100 μL, 0.5% in PBS) 1 hour before sacrifice. Tissue collected for Evans blue was assessed as previously described^83^; optical absorbance was measured with a Tecan Safire microplate reader, and the concentration of Evans blue was calculated^84^.

### Circulating tumor cell quantification

For circulating tumor cell (CTC) isolation, whole blood was collected at the day of sacrifice and centrifuged at 1200 rpm for 8 minutes at 4 °C. Pellet fraction was washed three times with ACK solution (BP10-548E, Lonza) for osmotic lysis of red blood cells and then seeded in complete medium. After 10 days, tumor colonies were fixed with cold methanol and stained with crystal violet, and then manually quantified.

### In vivo BLI

*In vivo* BLI was performed during the first 3–14 days for HIF-1 activity by exGLuc-based imaging of the primary tumors with a water-soluble Coelenterazine (NanoLight Technology, Pinetop, AZ, USA) (50 μg in 50 µl, i.v.) and for FLuc activity each week during experiments for metastatic development with 50 μl of D-Luciferin (30 mg/ml; i.p.) (Gold Biotechnology, St. Louis, USA). Mice were anesthetized prior to injection, and images were acquired using an IVIS Imaging system (WCM Animal Imaging/PET service). Photons emitted from the tumor region were quantified using Living Image software (PerkinElmer, Caliper Life Sciences, Mountain View, CA).

### [^18^F]FDG/PET

[^18^F]FDG PET/CT was acquired on WT mice injected with 4T1-KD and 4T1-Sc cells (1 × 10^6^) in contralateral m.f.p. on day −14 and −11, respectively, to normalize tumor volume at the time of readout. After being fasted overnight, mice were administered i.v. with 6.1 ± 0.1 MBq of [^18^F]FDG (Sofie Biosciences). A 10-minute static image was then acquired 1.5 hours post-injection on a Siemens Inveon PET/CT system. Calibrated cross-sectional and maximum intensity projection (MIP) renderings were prepared using Inveon Research Workspace (IRW; Seimens). A biodistribution was performed immediately following the PET/CT acquisition. The animals were euthanized using CO_2_ asphyxiation and heart puncture, select tissues were then excised, weighed and measured on a PerkinElmer Wizard^2^ 3” 2480 gamma counter on the 511 keV photopeak of F-18. A calibration factor was applied to convert the data from CPM to MBq, decay corrected to the time of injection and is presented as the percentage of injected dose per gram of tissue (% ID/g).

### DCE MRI

4T1-Sc and 4T1-KD tumors were implanted in contralateral m.f.p. of the same mouse to reduce variations in host vasculature impacting the assessment of the tumor vasculature. Prior to DCE MRI, mice were catheterized to facilitate the administration of the contrast agent (CA) Gd-DTPA (0.2 mmol Gd-DTPA/kg body weight, Magnevist®, Berlex Laboratories, Inc., Wayne, NJ) via a custom-built catheter line assembly. The catheter was kept patent by heparinized saline. The DCE-MRI experiments were performed on a 9.4T MR imager (BioSpec 94/20, Bruker, Germany), using a commercial cardiac ^1^H MR coil (4-loop phase array, Bruker, Germany) in receive mode inside a whole-body ^1^H MR coil (35 mm inner diameter, Bruker, Germany) for excitation. Mice were anesthetized with 1-3% isoflurane in 100% oxygen for the duration of the MR experiment. Mice were placed prone on the cardiac coil with the loops of the cardiac coil covering both tumors on the abdomen of the mouse. A respiration monitor pad (SA Instruments, Inc., Stonybrook, NY) was placed under the abdomen (above the cardiac coil) for respiratory monitoring during the MR experiment. The setup was then positioned in the center of the whole-body coil. The entire coil & mouse assembly was centered in the magnet, followed by tuning, matching of the coils to ^1^H frequency, and field-map-based shimming to eliminate (reduce) magnetic field inhomogeneities. During the MR experiments, the respiration rate was kept between 40-90 breaths / minute by adjusting the isoflurane concentration. For the MR acquisitions in one of the two cohorts, the mouse body temperature was controlled and kept at (36.01±0.25) °C (n = 8) during the DCE MRI scans using an MR-compatible small animal water heating system (Small Animal Instruments, Inc., Stony Brook, NY). After a pilot scan to determine the number of axial slices required to entirely cover both tumors, a spin-density MR image (Fast Low Angle Shot (FLASH) MRI, 500 ms repetition interval (TR), 1.346 ms echo time (TE), 1 repetition (NR), 1 average (NA), 1 mm slice thickness (st), 30 mm x 20 mm field of view (FOV), 192 x 128 matrix size, 6-12 slices; 15° flip angle) was acquired to visualize anatomy and facilitate tumor and dLN volume and region of interest outlining (V_DCE_) for subsequent DCE MRI data analysis. The DCE MRI data were acquired using FLASH with the same FOV, slices, st, matrix size, TE, NA, and flip angle as the spin density image, with TR set to the minimum TR permitted by the number of slices to maximize the temporal resolution. Gd-DTPA was injected via tail vein after 2 minutes of baseline acquisition followed by the dynamic acquisition.

DCE-MRI data were analyzed using principal component analysis (PCA) followed by constrained non-negative matrix factorization (cNMF), thus, estimating the spatial distributions of well-perfused (fast CA wash-in/washout, well oxygenated) tumor areas (P, pattern 1, P1), and tumor areas characterized by delayed CA wash-in/washout (typically associated with hypoxic tissue) (H, pattern 2, P2) and by slow or no accumulation of CA with no or marginal CA washout (no functional vasculature, typically associated with necrosis) (N, pattern 3, P3)^85–88^. Single pixel signal time curves were fitted using the Hoffman model^89^, to obtain the parameter Ak_ep_, a quantitative measure of blood flow & permeability. Median Ak_ep_ values were calculated for the whole tumor (Median Ak_ep_(whT)) and the tumor areas characterized by the three different CA uptake and washout behaviors (Ak_ep_(P), Ak_ep_(H), Ak_ep_(N)). We also calculated an Ak_ep_ value for the whole tumor based on the average signal-time curve (Average Curve Ak_ep_(whT)). Further, the volume fraction of tumor associated with pixels representing each of the 3 CA wash-in/-out behaviors (%V_P1_, %V_P2_, %V_P3_) and with pixels characterized by more than one behavior (%V_P1+P2_, %V_P1+P3_, %V_P2+P3_, %V_P1+P2+P3_) were obtained from the pattern recognition analysis, as before^90,91^.

### MALDI Imaging Metabolomics

Tumors were surgically resected to preserve metabolites and freshly frozen in isopentane before embedding in 2% carboxymethylcellulose. Embedded samples were stored at −80°C for at least 24-48 hours and placed at −20°C the day before sectioning. Tissue sections (10 µm) were placed on the conductive side of indium-tin-oxide (ITO) coated glass microscope slides and acquired for untargeted metabolites at 40µm spatial resolution using NEDC as a matrix and negative ion mass spectrometry detection by 7T ScimaX (Bruker Daltonics, USA) magnetic resonance imaging mass Spectrometer (MRMS) at the Weill Cornell Advanced Biomolecular Analysis Core Facility. Pixel-by-pixel tissue intensity for each metabolite was extracted as a measure of metabolite abundance.

### Analyses of transcriptomic data

The murine data set was obtained from Zappasodi et al.^13^. Human data sets from TCGA^41^ were obtained through the UCSC Xena platform^92^. The gene signatures for HEVs and tumor ECs were obtained from Hua et al.^39^. For the analysis of the mouse data set, the original signatures from the study were used: the TU EC signature for tumor ECs, and a combined HEV signature built by merging genes from the TU HEV and the HEV common signatures. For the human data sets, we used the human TU HEV gene signature from the same source. The signature for tumor vasculature was obtained from Lu et al.^93^ as part of the HALLMARK angiogenesis signature. Signatures representing angiogenic and quiescent PCs were obtained from Teuwen et al.^35^, while the signature for cytolytic activity was derived from Rooney et al.^40^. Signatures for metabolic pathways were constructed using a knowledge-based approach. For signatures lacking a direct human–mouse counterpart, orthologous genes were identified via BioMart^94^, and the corresponding signatures were then constructed accordingly. A complete list of genes included in each signature is provided in **Suppl. Table 1**. The combined signature was derived using the arithmetic mean of glycolysis, HEV, tumor EC, quiescent PC, angiogenic PC, and cytolytic activity standardized scores. Scores for signatures associated with a protective effect (HEV, cytolytic activity, and quiescent PCs) were sign-inverted prior to standardization. To calculate the per-sample score for each signature, the GSVA method was applied using the GSVA R package, utilizing the Gaussian kernel cumulative distribution function. To estimate the immune cell fractions from bulk RNA-seq data, we performed deconvolution using the MCPcounter method^95^ for human data and mMCPcounter^96^ for the murine data set, both implemented through the immunedeconv R package. Transcriptomic data and related progression free survival (PFS) information from HCC patients treated with atezolizumab vs. atezolizumab with bevacizumab in group F of the GO30140 phase I trial investigating atezolizumab vs. atezolizumab plus bevacizumab were obtained from EGAS00001005503^38^. Tumor samples were divided into glycolysis-high and -low based on the median glycolysis signature score (calculated as above) and PFS for patients treated with atezolizumab vs. atezolizumab plus bevacizumab was compared within each tumor glycolysis condition. Hazard ratios were calculated using a Cox proportional regression model.

### Statistical analyses

The normality of data distributions was evaluated using the Shapiro-Wilk test. Parametric (unpaired *t* test and one-way or two-way ANOVA followed by Bonferroni’s correction for multiple comparisons) or non-parametric (Mann-Whitney U test and one-way Kruskal-Wallis H test followed by Dunn’s post-hoc statistical analysis) was used. P values for survival analyses were calculated using the log-rank (Mantel-Cox) test. Statistical analyses were performed using GraphPad Prism 10.5 software.

Statistical significance for the vasculature signatures scores between treatment conditions using the murine data set was assessed using two-sided pairwise Wilcoxon rank-sum tests. To quantify the correlation between signature scores, Pearson correlation coefficients, and their relative significance were calculated using the rcorr function from the Hmisc R package. Clinical data for every TCGA sample were retrieved from Liu et al.^97^. Signature scores were z-standardized with the StandardScaler class from the scikit-learn library. To assess the association between each signature and PFI, a univariate Cox proportional hazards model was fitted for each one of the signatures and the combined signature for every tumor type in the TCGA data set using the class CoxPHFitter from the lifelines Python package. The lifelines library was also employed to generate Kaplan–Meier survival curves with the KaplanMeierFitter class.

Detailed information of the statistical test and number of observations/replicates used in each experiment, and the definition of center and dispersion is appropriately reported in the legend of each figure. Significance was defined as follows: * = p<0.05, ** = p<0.01, *** = p<0.001, **** = p<0.0001.

## Acknowledgements

We thank the Advanced Biomolecular Analysis Core Facility and Flow Cytometry Core Facility at Weill Cornell Medicine, as well as the Molecular Cytology Core Facility and MRI Facility at Memorial Sloan Kettering (supported by Core Grant P30 CA008748), for their technical assistance. We thank the NIH Tetramer Core Facility (NIH Contract 75N93020D00005 and RRID:SCR_026557) for providing the H-2Db/gp100_25–33 tetramer.

This research was supported in part through the Bristol Myers Squibb International Immune Oncology Network, the Breast Cancer Alliance Young Investigator grant, the Cancer Research Institute CLIP Grant (CRI Award #13928), and the G. Harold and Leila Y. Mathers Charitable Foundation (R.Z.); and by R01A CA215136 (R.B.). I.S. was supported by R50 CA221810 (NIH/NCI) grant. G.C. was supported by the AIRC abroad postdoc fellowship (Project Code: 29682). F.B. was supported by European Union—NextGenerationEU: National Center for Gene Therapy and Drugs based on RNA Technology, CN3—Spoke 7 (code: CN00000041; PNRR—Mission 4, Component 2; Investment 1.4). J-H.K. was supported by the Parker Institute for Cancer Immunotherapy RISE Scholar Award. T.V.F.E. was supported by the Tow Post Doctoral Fellowship. A.Santella was supported by the Chan Zuckerberg Initiative Imaging Scientist Grant. E.Andreopoulou was supported by the Kat’s Ribbon of Hope Organization. R.Z. is the recipient of the Emilie Lippmann and Janice Jacobs McCarthy Research Scholar in Breast Cancer Award at Weill Cornell Medicine.

## Author contributions

I.S., G.C., R.Z. developed the concept, designed and discussed experiments; G.C., I.S., R.Z. wrote the manuscript; I.S., G.C., J.H.K, E. Ackerstaff, T.K., T.V.F.E. performed and analyzed experiments; F.B., P-F.G., A.Santella performed bioinformatic analyses; A.Schreier, E.Andreopoulou, S.D. provided human tissue and contributed to human tissue analyses; A.W.L. provided Kaede mice and scientific input; N-V-K. P. provided scientific input; R.B., R.Z. secured research funding; R.Z., I.S. supervised the research.

