## Supplementary Files for "A tumor metabolism-angiogenesis-immune axis governs immunotherapy responses"

### Supplemental Figures

#### Supplemental Figure 1

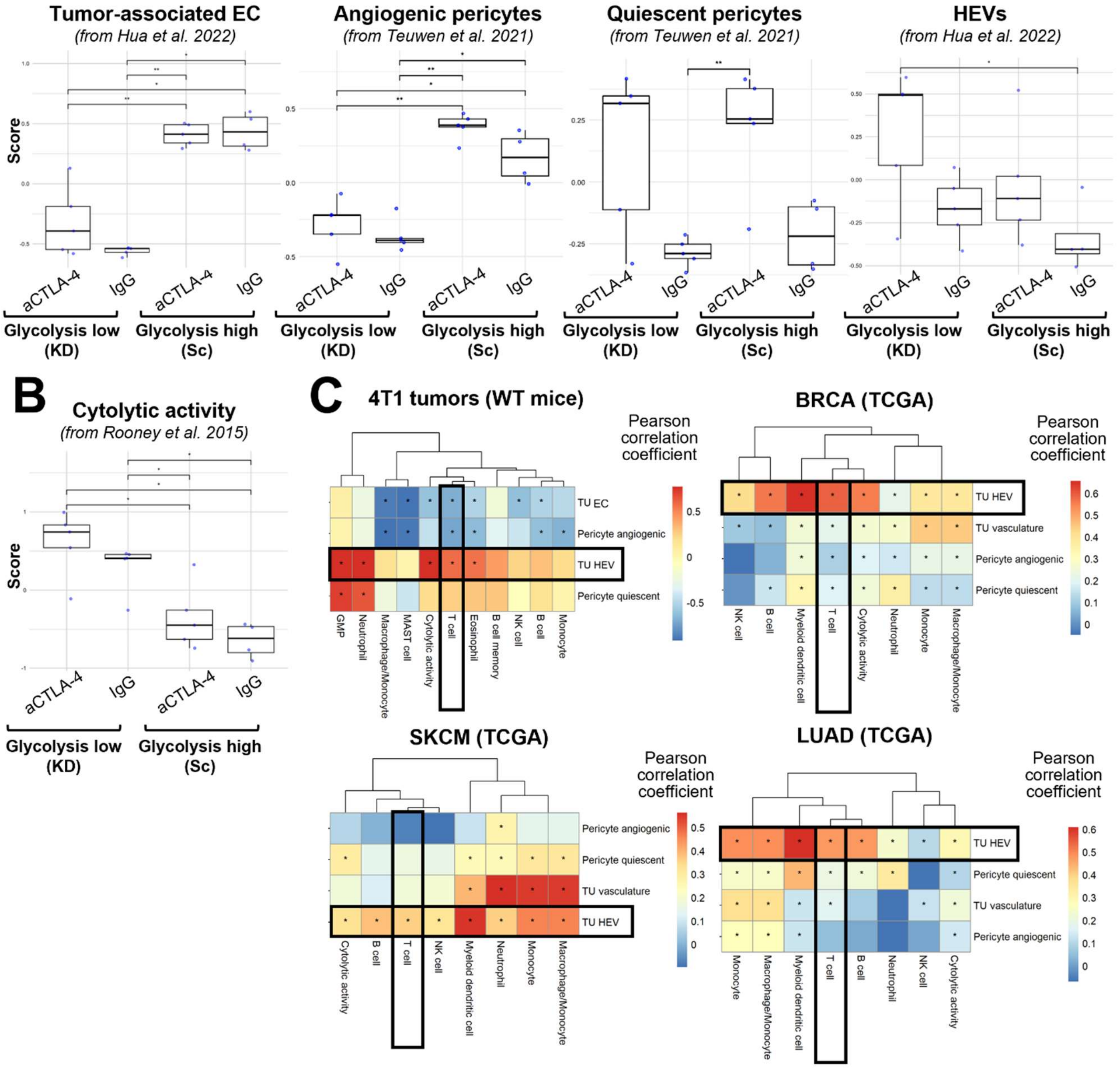

**Supplemental Figure 1: Relationship between tumor cell glycolysis, tumor vasculature features and immunotherapy responses (related to Fig. 1).** Enrichment scores for the indicated signatures representing tumor-associated ECs, angiogenic PCs, stabilizing quiescent PCs, and HEVs (**A**) and (**B**) cytolytic activity in RNAseq data set from LDHA-KD and Sc 4T1 tumors treated with IgG or anti-CTLA-4 (9D9) (GSE164051). Statistical significance was calculated by 2-sided pairwise Wilcoxon rank-sum tests (*, p<0.05; **, p<0.01). **(C)** Enrichment scores of vasculature signatures (Suppl. Table 1) were correlated with immune infiltrating cells estimated by standard deconvolution methods using mMCPcounter (for mouse RNAseq data) and MCPcounter (for human TCGA data sets), and cytolytic activity estimated according to Rooney et al. Cell 2015. *, significant results by Pearson correlation.

**Supplemental Figure 2**

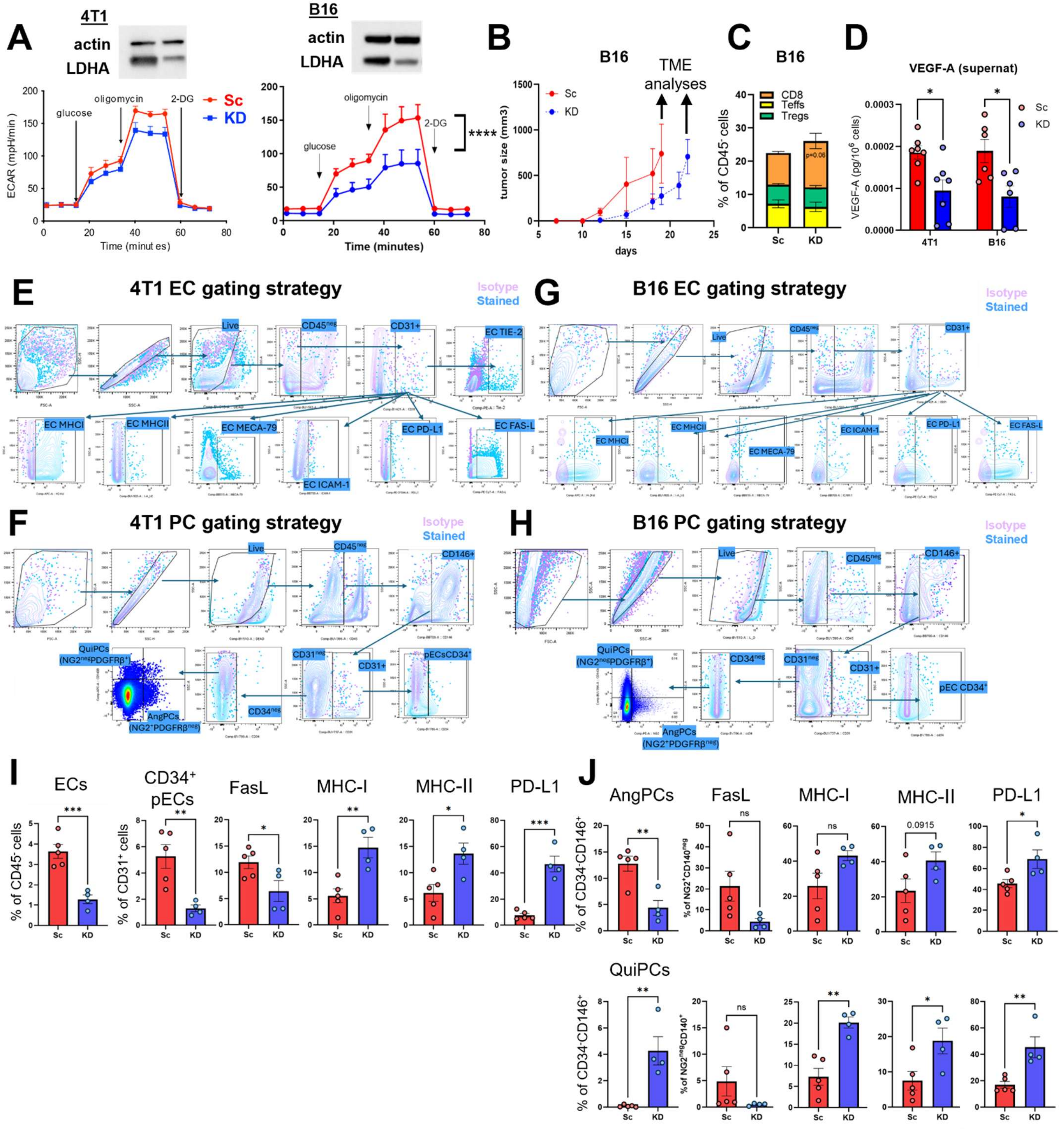

**Supplemental Figure 2: Tumor LDHA-KD reshapes the tumor vasculature phenotype (related to Fig. 2).** (**A**) Western blot protein expression analysis of LDHA and actin as loading control and extracellular acidification rate (ECAR) by Seahorse assay comparing LDHA-KD and Sc variants of 4T1 and B16 tumor cells. (**B**) Growth curves of B16-Sc and B16-KD in WT C57BL/6 mice (n=5/group). Arrows indicate tumor harvesting time points for flow cytometry analyses of the tumor vasculature and immune infiltrate when primary tumors reached similar sizes in (C, G-J). (**C**) Flow cytometry quantification of the indicated intratumor T-cell subsets ($CD8^+$, $Foxp3^-CD4^+$ Teff, $Foxp3^+CD4^+$ Treg cells) in B16-Sc and B16-KD tumors harvested from WT mice as in (C). (**D**) Luminex beads-based immunoassay quantification of VEGF-A production in LDHA-KD and Sc 4T1 and B16 cell culture supernatants after 24-hr incubation. (**E-H**) Representative flow cytometry gating strategy for quantification of EC and PC subsets and their phenotype in 4T1 and B16 tumors (isotype, cells were stained for lineage markers and viability dye and with fluorochrome-conjugated isotype controls corresponding to the antibodies used in the fully stained samples). (**I, J**) Flow cytometry quantification of frequencies and expression of the indicated markers in ECs (I), and (J) angiogenic PCs (Ang PCs) and quiescent PCs (QuiPCs) in B16-Sc and B16-KD tumors from WT mice as in (B). Median fluorescence intensity, MFI. Data are mean ± SE of one representative out of 3 independent experiments (n=4-5). *, $P<0.05$; **, $P<0.01$; ***$P<0.001$.

**Supplemental Figure 3**

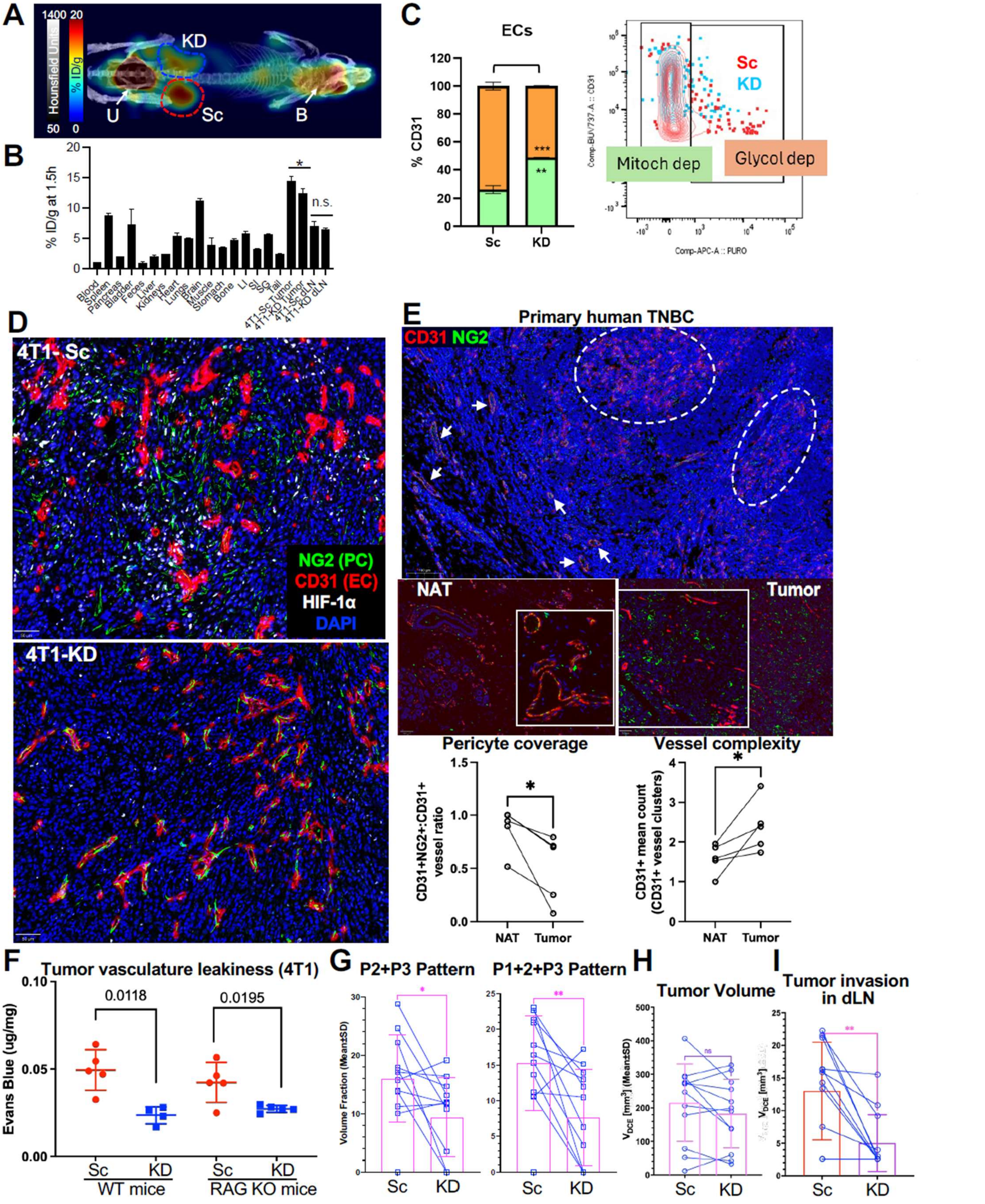

**Supplemental Figure 3: Tumor vasculature normalization in glycolysis-defective vs. glycolytic tumors (related to Fig. 3).** (**A**) Representative maximum intensity projection (MIP) PET/CT of [$^{18}$F]FDG in mice bearing contralateral 4T1-KD and 4T1-Sc tumors in the m.f.p. at 1.5 h post tracer injection as in Fig. 3A. B, brain; U, urine within bladder (PET signal in colormap, CT in greyscale). (**B**) Complete biodistribution of [$^{18}$F]FDG in mice bearing contralateral 4T1-KD and 4T1-Sc tumors at 1.5 h post tracer injection as in Fig. 3A (n=4; mean +/- SE; 2-sided unpaired t test. *, P<0.05). (**C**) Metabolic analysis of EC isolated from Sc and KD tumors by SCENITH, showing quantification of glycolytic (orange) vs. mitochondrial (green) dependence and related representative flow cytometry plots (puromycin incorporation was used as a readout for ATP production under oligomycin treatment, where puromycin$^{+}$ cells indicate glycolytic dependence and puromycin$^{-}$ cells indicate mitochondrial dependence) (n=8/group). (**D**) Representative multiplexed immunofluorescence (IF) staining for NG2 PC, CD31 EC, and HIF-1$\alpha$ hypoxia markers in 4T1-Sc and 4T1-KD tumors (quantified in Fig. 3E-H). (**E**) Representative images and quantification of IF staining for NG2 and CD31 in human primary TNBC cases comparing vasculature structures by PC coverage (ratio between CD31$^{+}$NG2$^{+}$ vessel structures over total CD31$^{+}$ vessel structures) and complexity (mean number of CD31$^{+}$ ECs in vessel structures) in tumor core (dotted circles) vs. or normal adjacent tissue (NAT; arrows) (n=5). (**F**) Tumor vasculature leakiness by Evans blue assay in 4T1-Sc and 4T1-KD tumors grown in WT vs. RAG2 KO mice (n=4-5). (**G**) DCE-MRI-based quantification of % tumor volume fractions displaying mixture patterns indicative of reduced perfusion (P1+P2, left; and P1+P2+P3, right) overlaid with corresponding mean ±SD (n= 12) in 4T1-Sc (Sc) and 4T1-KD (KD) tumors implanted in contralateral m.f.p. of the same mouse (showing individual data that were averaged and reported in Fig. 3K). (**H**) Quantification of tumor volume by MRI in WT mice bearing contralateral 4T1-Sc (Sc) and 4T1-KD (KD) tumors as in (F). (**I**) Quantification of dLN volume by MRI showing significantly more (9/12) invaded and larger ilioinguinal dLN for 4T1-Sc (Sc) than similar-sized 4T1-KD (KD) tumors (5/12) from same mice as in (G, H). (**G-I**) Data are from two independent cohorts combined in which tumors were harvested when they reached ~12-130 mm$^3$ (n=3) or ~150-440 mm$^3$ (n=9). Pairwise comparison shows no difference in tumor size between 4T1-Sc and 4T1-KD at the time of MRI acquisition. 2-sided paired (E, G-I) or unpaired (B, C, F) *t* test: *, P<0.05; **, P<0.01; ***, P<0.001.

**Supplemental Figure 4**

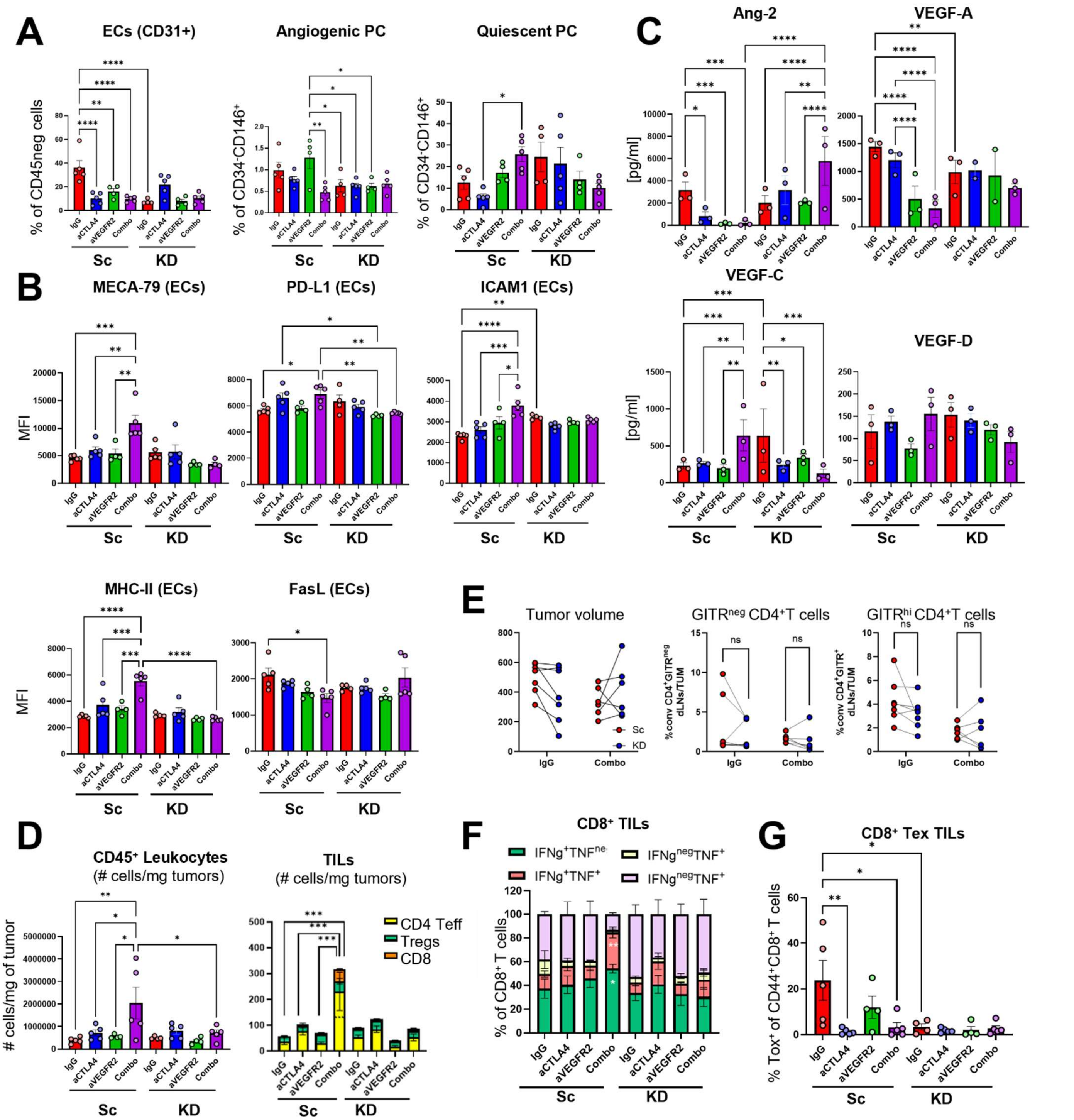

**Supplemental Figure 4: Anti-VEGFR2 plus anti-CTLA-4 improves the TME of glycolytic but not glycolysis-defective tumors (related to Fig. 5).** (**A-D**) B16-Sc an B16-KD tumors from mice treated with anti-CTLA-4 $\pm$ anti-VEGFR2 or isotype control as in Fig. 5B were processed for: (A) flow cytometry quantification of frequencies of the indicated vascular cells; (B) expression (by MFI) of the indicated markers in ECs; (**C**) TIF analyses of the indicated hem/lymphangiogenic factors quantified by Luminex bead-based immunoassays; (**D**) flow cytometry quantification of $CD45^+$ leucocytes and tumor infiltrating lymphocytes (TILs: $CD8^+$, $CD4^+Foxp3^-$ Teff, $CD4^+Foxp3^+$Tregs) per mg of tumor. (**E**) Tumor volume and flow cytometry quantification of ratio photoconverted $CD4^+GITR^-$ T cells, $CD4^+GITR^{hi}$ Tregs (discriminated by surface staining of GITR for compatibility with Kaede protein detection) in dLN relative to tumor (TUM) in control vs. combo-treated B16-Sc and B16-KD Kaede mice as in Fig. 5L. (**F,G**) Frequencies of $CD8^+$ T cells expressing IFN-$\gamma$ $\pm$ TNF (F), and (G) frequencies of antigen-experienced $CD44^+CD8^+$ T cells expressing the exhaustion marker Tox by flow cytometry in the same tumors as in (A-D) treated as in Fig. 5B. Data are mean $\pm$ SE (n=3-5/group). *, P<0.05; **, P<0.01; ***, P<0.001; ****, P<0.0001.

**Supplemental Figure 5**

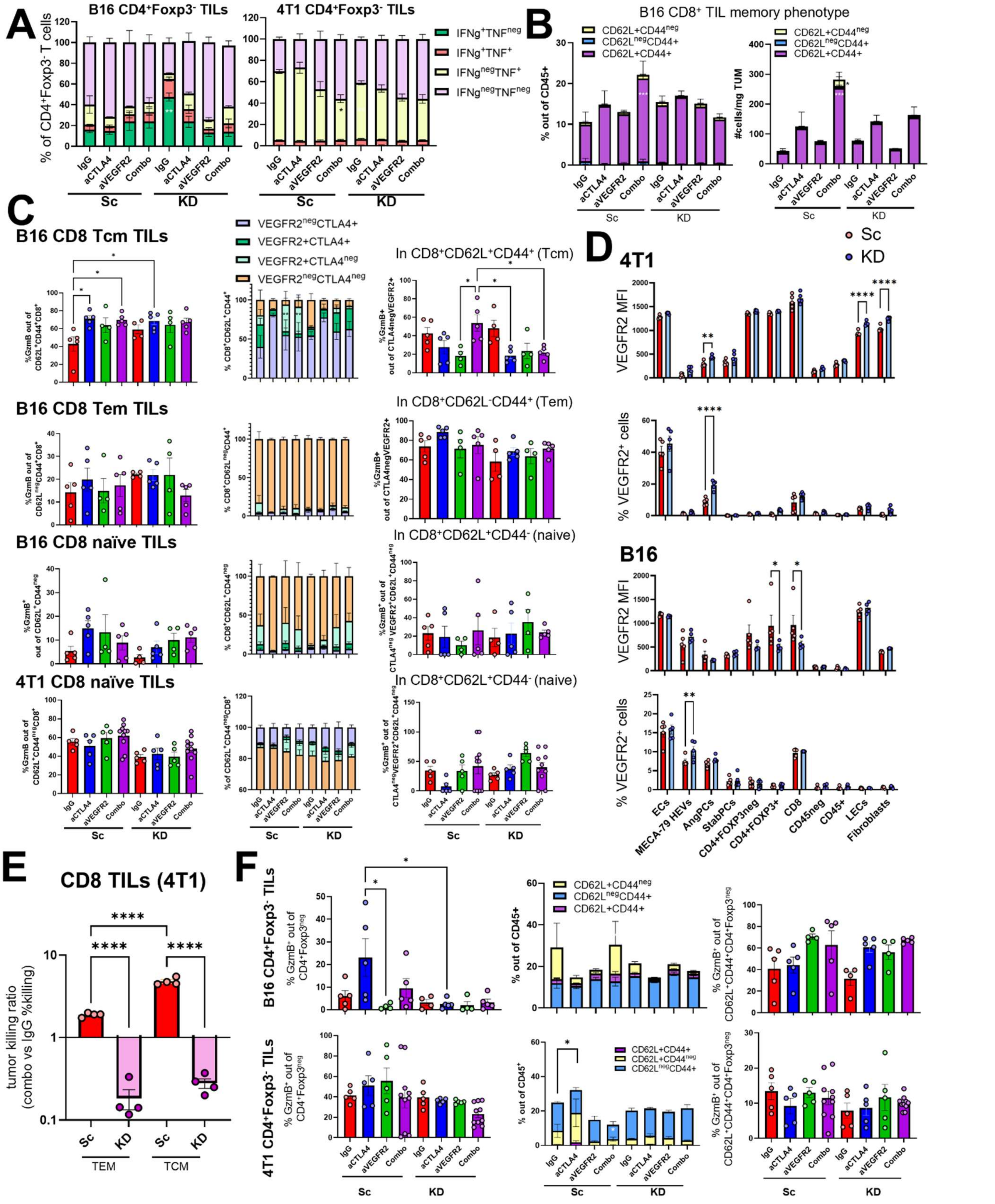

**Supplemental Figure 5: Intratumor T-cell functional changes upon anti-CTLA-4 plus anti-VEGFR2 treatment according to the tumor glycolytic state. (related to Fig. 6).** (**A**) Flow cytometry quantification of frequencies of CD4$^+$Foxp3$^-$ T cells expressing IFN-$\gamma$ $\pm$ TNF in B16-Sc vs B16-KD (left) and 4T1-Sc vs 4T1-KD (right) from mice treated as in Fig. 5A,B. (**B**) Flow cytometry quantification of CD8$^+$ CD62L$^+$CD44$^-$ naïve, CD62L$^+$CD44$^+$ central memory (Tcm), and CD62L$^-$CD44$^+$ effector memory (Tem) cell frequencies and absolute numbers per mg of tumor in B16-Sc and B16-KD tumors treated as in Fig. 5B. (**C**) Flow cytometry quantification of GzmB$^{+}$ cells (left), VEGFR2$^{+/-}$CTLA-4$^{+/-}$ cell fractions (middle), and (right) VEGFR2$^{+}$CTLA-4$^{-}$ cells expressing GzmB within CD8$^{+}$ Tcm, Tem and naïve cells from B16 and/or 4T1 tumors treated as in Fig. 5A,B. (**D**) Flow cytometry quantification of VEGFR2 expression (as % positive cells and MFI) in the indicated stromal cells in LDHA-KD and Sc 4T1 (top) and B16 tumors (bottom) from WT mice. (**E**) Ratio % tumor killing by FACS-sorted CD8 Tcm (CD8$^{+}$CD62L$^{+}$CD44$^{+}$) and Tem (CD8$^{+}$CD44$^{+}$CD62L$^{neg}$) from combo-treated vs. IgG-treated 4T1-Sc and 4T1-KD tumors (1 representative of 3 independent experiments with either 4T1 or B16 models). (**F**) Flow cytometry characterization of tumor-infiltrating CD4$^{+}$Foxp3$^{-}$ T cells from LDHA-KD and Sc B16 (top) and 4T1 (bottom) tumors treated with anti-CTLA-4 4 $\pm$ anti-VEGFR2 or isotype control as in Fig. 5A,B: % cells expressing GzmB (left), CD44 $\pm$ CD62L (middle), and (right) GzmB within the Tcm subset. Data are mean $\pm$ SE (n=5-10/group in one of 2-3 independent experiments). *, P<0.05; **, P<0.01; ***, P<0.001; ****, P<0.0001.

**Supplemental Figure 6**

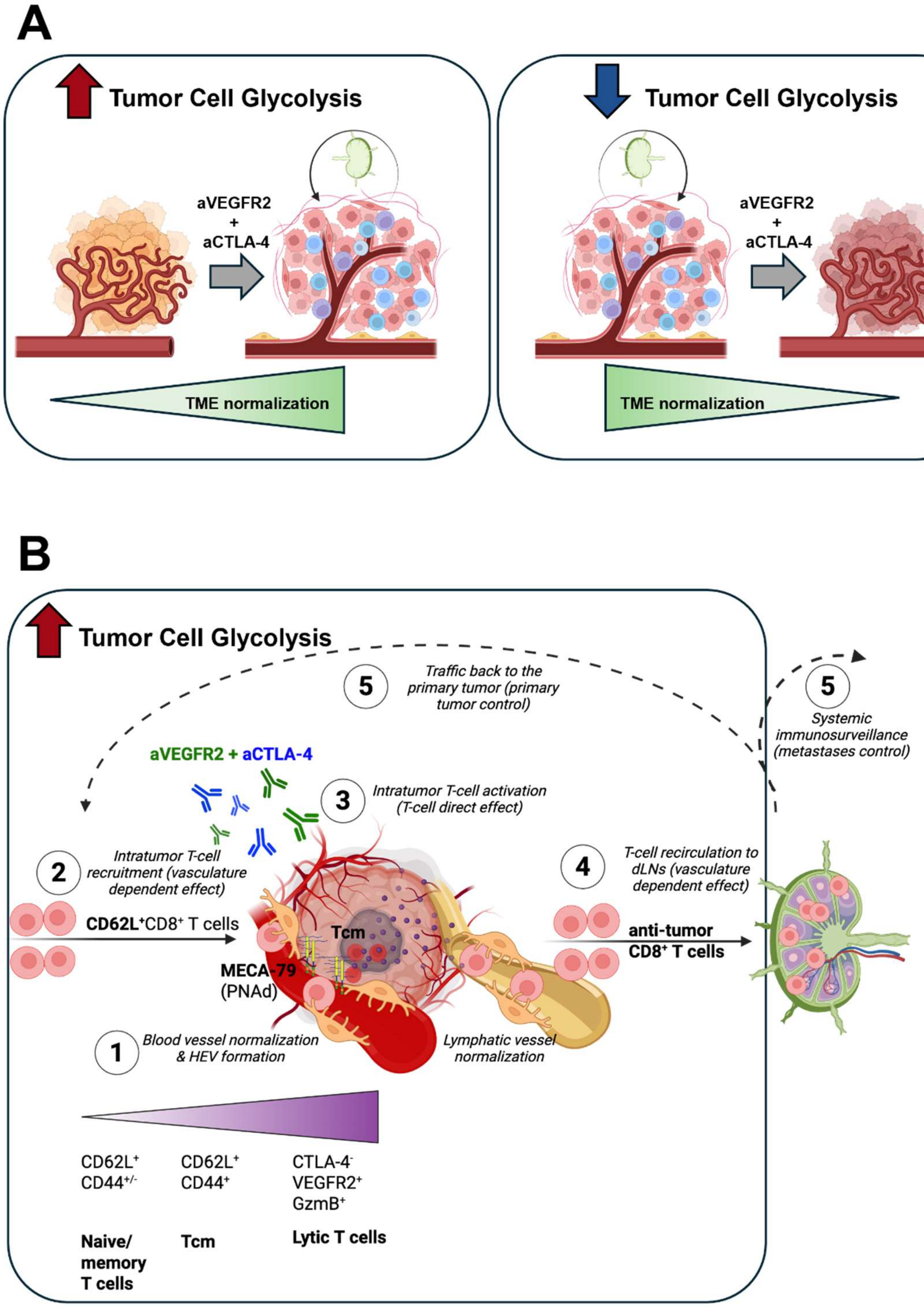

**Supplemental Figure 6: Models summarizing findings. (A)** Schematic representation of how the tumor glycolytic state impacts the TME immune and vascular normalization upon treatment with anti-CTLA-4 plus anti-VEGFR2. (**B**) Schematic representation of the sequence of events triggered by anti-VEGFR2 plus anti-CTLA-4 in glycolytic tumors supporting vasculature normalization and immune responses.

### Supplemental Tables

#### Supplemental Table 1: Gene signatures used in the study.

| Mitochondrial | Carbon metabolism | FA synthesis | Glycolysis | FA Catabolism | PC quiescent | PC angiogenic | Cytolytic activity | TU HEV | TU vasculature |
|---|---|---|---|---|---|---|---|---|---|
| mt-Co2 | Cbs | Elovl5 | Pfkl | Slc27a1 | Ltbp2 | Fstl1 | Gzma | Glycam1 | Nid2 |
| mt-Co1 | Mtrr | Elovl7 | Pgk2 | Slc27a2 | Klf9 | Acta2 | Prf1 | Chst4 | Flt4 |
| mt-Nd6 | Mat1a | Scd2 | Pgam1 | Hadha | Fos | Col1a2 | | Serpina1b | Lama4 |
| mt-Nd2 | Psat1 | Scd4 | Gpi1 | Acsl4 | Itm2a | Col3a1 | | Enpp2 | Sema6d |
| mt-Atp8 | Tyms | Elovl4 | Pfkm | Hadhb | Cystm1 | Col5a2 | | Apoe | Dbn1 |
| mt-Cytb | Bhmt | Acaca | Hk2 | Cpt2 | Cryab | Serpine2 | | C1s1 | Piezo2 |
| mt-Nd4 | Dhfr | Slc25a1 | Aldoa | Acsl6 | Pdzd2 | Basp1 | | Gpr182 | Emid1 |
| mt-Nd5 | Shmt2 | Elovl3 | Aldoart2 | Ehhadh | Cox4i2 | Serpina1b | | Clu | Kcne3 |
| mt-Co3 | Fpgs | Fads1 | Aldoart1 | Ppa1 | Prex2 | Actg2 | | Ubd | Nes |
| mt-Atp6 | Folh1 | Elovl6 | Pgk1 | Slc25a20 | Nkain4 | Thy1 | | Lifr | Tspan14 |
| mt-Nd1 | Gldc | Fasn | Slc16a3 | Acadl | Npnt | Anxa1 | | Cfb | Plod1 |
| mt-Nd4l | Psph | Acly | Bsg | Echs1 | Enpp2 | Fabp5 | | Spint2 | S100a6 |
| mt-Nd3 | Gnmt | Fads2 | Eno2 | Acsl5 | Tsc22d1 | Tm4sf1 | | Ctsl | Mest |
| mt-Rnr1 | Ggh | Elovl2 | Tpi1 | Acsf3 | Mfge8 | Vim | | Il2rg | Eogt |
| mt-Rnr2 | Ahcy | 9130409I23Rik | Pklr | Acat1 | Dusp1 | Rps25 | | Gm10851 | Ecm1 |
| mt-Tk | Mthfd1l | Degs1 | Ldhb | Cpt1a | Rgcc | Rpl41 | | Sult1a1 | Notch4 |
| mt-Tr | Mthfs | Aacs | Pgam2 | Acadvl | Hbegf | Mmp14 | | Serpinb9 | Bmp1 |
| mt-Ta | Mthfd1 | Elovl1 | Pfkp | Acsl3 | Crip2 | Serf2 | | Tmem176a | Gpx3 |
| mt-Ty | Cth | | Pkm | Acsf2 | Ckb | Mgp | | Lrg1 | Slc44a2 |
| mt-Tw | Mthfd2l | | Hk1 | Acads | Lmcd1 | Cd248 | | Fth1 | Cd276 |
| mt-Tv | Mthfd2 | | Gm10358 | Acsl1 | Malat1 | Gm8730 | | Il6st | Podxl |
| mt-Tt | Phgdh | | Gm3839 | Acsm4 | G0s2 | Eef1g | | C1ra | Plat |
| mt-Ts2 | Chdh | | Gapdh | Acadm | Tppp3 | Col6a3 | | Ackr1 | Ramp3 |
| mt-Ts1 | Mtr | | Eno3 | | Higd1b | Rpl3 | | Csf2rb2 | Tubb2a |
| mt-Tq | Shmt1 | | Slc16a1 | | Ccrl2 | Timp1 | | Tmem176b | Fn1 |
| mt-Tp | Mthfr | | Ldha | | Gpx3 | Col18a1 | | Abcg2 | Sipa1 |
| mt-Tn | | | Slc2a1 | | Sftpc | Serpinh1 | | Csf2rb | Tspan18 |
| mt-Tm | | | Eno1 | | Ecm1 | Rpl32 | | B2m | Anxa6 |
| mt-Tl2 | | | Eno1b | | Shroom3 | Lgals1 | | Osmr | Abcc9 |
| mt-Tl1 | | | | | Hspa1a | Arf4 | | H2-K1 | Apln |
| mt-Ti | | | | | Klf2 | Rplp0 | | Cept1 | Col13a1 |
| mt-Th | | | | | Tbx5 | Rgs5 | | Xbp1 | Afap1l1 |
| mt-Tg | | | | | Pde5a | Tpm1 | | Cxcl1 | Kcnq1 |
| mt-Tf | | | | | Clec3b | Fn1 | | Pim3 | Tmem204 |
| mt-Te | | | | | Mxra8 | Col15a1 | | Ehd3 | Ptn |
| mt-Td | | | | | Rarres2 | Rps12 | | Serpinb1a | Ackr3 |
| mt-Tc | | | | | Ism1 | Gapdh | | Clec14a | Mmp14 |
| | | | | | Ifitm1 | Rpl38 | | Cd300lg | Dll4 |

| | | | | | | | | | |
|---|---|---|---|---|---|---|---|---|---|
| | | | | | Prelp | Rpl18 | | Egr1 | Lamc1 |
| | | | | | Zeb2 | Col1a1 | | Cd36 | Esm1 |
| | | | | | Cdkn1c | Rpsa | | Ifitm3 | Cldn5 |
| | | | | | Ppp1r14a | Igfbp7 | | Selp | Peg3 |
| | | | | | Aldh2 | Rps19 | | Mctp1 | Ptp4a3 |
| | | | | | Kcnk3 | | | Ptafr | Igfbp5 |
| | | | | | Tbx2 | | | Tm4sf1 | Gja4 |
| | | | | | Gsn | | | Tspan7 | |
| | | | | | Cst3 | | | Sele | |
| | | | | | | | | Gm37376 | |
| | | | | | | | | Slc39a1 | |
| | | | | | | | | Iigp1 | |
| | | | | | | | | Ifitm2 | |
| | | | | | | | | Car4 | |
| | | | | | | | | Gda | |
| | | | | | | | | Il6 | |
| | | | | | | | | Slco3a1 | |
| | | | | | | | | Vwf | |
| | | | | | | | | Meox2 | |
| | | | | | | | | H2-Q6 | |
| | | | | | | | | Adrb2 | |
| | | | | | | | | Pdlim1 | |
| | | | | | | | | Casp4 | |
| | | | | | | | | Gbp9 | |
| | | | | | | | | Tmem252 | |
| | | | | | | | | Vcam1 | |
| | | | | | | | | Il1r1 | |
| | | | | | | | | Fabp5 | |
| | | | | | | | | Cebpd | |
| | | | | | | | | Fbln2 | |
| | | | | | | | | Ablim1 | |
| | | | | | | | | Igtp | |
| | | | | | | | | Hdc | |
| | | | | | | | | Cxcl10 | |
| | | | | | | | | Lepr | |
| | | | | | | | | Rnd1 | |
| | | | | | | | | Dpysl3 | |
| | | | | | | | | Ctla2a | |
| | | | | | | | | Mustn1 | |
| | | | | | | | | Gadd45g | |
| | | | | | | | | Dnm3 | |
| | | | | | | | | Fosl2 | |
| | | | | | | | | Tgtp1 | |
| | | | | | | | | Clca3a2 | |
| | | | | | | | | Hspb1 | |
| | | | | | | | | Noct | |
| | | | | | | | | Rcan1 | |

| | | | | | | | | |
|---|---|---|---|---|---|---|---|---|
| | | | | | | | | Dst |
| | | | | | | | | Tap1 |
| | | | | | | | | Adamts9 |
| | | | | | | | | Ripk3 |
| | | | | | | | | Socs2 |
| | | | | | | | | Gbp7 |
| | | | | | | | | Cav1 |
| | | | | | | | | Gbp4 |
| | | | | | | | | Cyp4b1 |
| | | | | | | | | Tgtp2 |
| | | | | | | | | Ier3 |
| | | | | | | | | Csf3 |
| | | | | | | | | Socs3 |
| | | | | | | | | Tinagl1 |
| | | | | | | | | Myc |
| | | | | | | | | Zfp36 |
| | | | | | | | | Ifi47 |
| | | | | | | | | Nfkbia |
| | | | | | | | | Nfkbiz |
| | | | | | | | | Serpina3g |
| | | | | | | | | Mgll |
| | | | | | | | | Icam1 |
| | | | | | | | | Anxa1 |
| | | | | | | | | Irf1 |
| | | | | | | | | Tnfaip3 |
| | | | | | | | | Irgm1 |
| | | | | | | | | Btg2 |
| | | | | | | | | Lpl |
| | | | | | | | | Cd74 |

**Supplemental Table 2: Antibodies used for flow cytometry staining.**

| Marker | Fluorochrome | Clone | Company | Catalog number | Dilution |
|---|---|---|---|---|---|
| CD45 | PerCP-Cy5.5 | 30-F11 | BioLegend | 103132 | 1:200 |
| CD45 | APC-Cy7 | 30-F11 | BD Biosciences | 559864 | 1:200 |
| CD45 | BUV396 | 30-F11 | BD Biosciences | 564279 | 1:200 |
| CD45 | AF700 | 30-F11 | BioLegend | 103128 | 1:200 |
| CD8 | APC-Cy7 | 53-6.7 | BD Biosciences | 557654 | 1:200 |
| CD8 | BUV563 | 53-6.7 | ThermoFisher | 365-0081-82 | 1:200 |
| CD8 | BB700 | 53-6.7 | BD Biosciences | 566409 | 1:200 |
| CD8 | BUV737 | 53-6.7 | BD Biosciences | 612759 | 1:200 |
| CD4 | BV650 | RM4-5 | BioLegend | 100555 | 1:200 |
| CD4 | BV421 | GK1.5 | BD Biosciences | 562891 | 1:200 |
| CD4 | BUV395 | RM4-5 | BD Biosciences | 740208 | 1:200 |
| FOXP3 | PE | FJK-16s | ThermoFisher | 11-5773-82 | 1:200 |
| MECA-79 | FITC | MECA-79 | ThermoFisher | 53-6036-82 | 1:150 |
| PD-L1 | AF700 | MIH5 | ThermoFisher | M036T03R03-A | 1:200 |
| PD-L1 | PECy5 | 10F.9G2 | BioLegend | 124344 | 1:200 |
| PD-L1 | PECy7 | MIH5 | ThermoFisher | 25-5982-82 | 1:200 |
| PD-L1 | BV711 | MIH5 | BD Biosciences | 563369 | 1:200 |
| ICAM-1 | PerCP-Cy5.5 | YN1/1.7.4 | BioLegend | 116124 | 1:200 |
| FAS-L | PECy7 | MFL3 | ThermoFisher | 25-5911-82 | 1:200 |
| Tie-2 | APC | TEK4 | BioLegend | 124010 | 1:50 |
| CD31 | BUV737 | 390 | BD Biosciences | 741740 | 1:200 |
| CD31 | BV421 | 390 | BD Biosciences | 563356 | 1:200 |
| CD34 | BV786 | RAM34 | BD Biosciences | 742971 | 1:100 |
| CD146 | BV421 | ME-9F1 | BD Biosciences | 740095 | 1:100 |
| CD146 | BB700 | ME-9F1 | BD Biosciences | 742280 | 1:100 |
| CD140b | BUV395 | APB5 | BD Biosciences | 740269 | 1:200 |
| CD140b | APC | APB5 | ThermoFisher | 17-1402-82 | 1:100 |
| NG2 | PE | REA989 | Miltenyi | 130-116-376 | 1:25 |
| H-2dB | BUV563 | AF6-88.5 | BD Biosciences | 748823 | 1:200 |
| H-2kD | BUV563 | SF1-1.1 | BD Biosciences | 748357 | 1:200 |
| MHC-II | BUV805 | M5/114.15.2 | BD Biosciences | 748844 | 1:200 |
| MHC-II | AF700 | M5/114.15.2 | Biolegend | 107622 | 1:200 |
| VEGFR2 | FITC | REA1116 | Miltenyi | 130-119-432 | 1:50 |
| VEGFR2 | PE-Vio® 770 | REA1116 | Miltenyi | 130-119-436 | 1:50 |
| IFN | PE-CF594 | XMG1.2 | BD Biosciences | 562303 | 1:100 |
| TNF | APC | MP6-XT22 | BD Biosciences | 554420 | 1:100 |

| | | | | | |
|---|---|---|---|---|---|
| TOX | PE | TXRX10 | ThermoFisher | 12-6502-82 | 1:40 |
| CD44 | BUV737 | IM7 | BD Biosciences | 612799 | 1:200 |
| CD44 | BV786 | IM7 | BioLegend | 103059 | 1:200 |
| CD44 | BV421 | IM7 | BioLegend | 103040 | 1:200 |
| CD62L | BV605 | MEL-14 | BioLegend | 104438 | 1:200 |
| CD62L | AF700 | MEL-14 | BD Biosciences | 560517 | 1:200 |
| CTLA4 | PeCy5 | UC10-4B9 | BioLegend | 106338 | 1:200 |
| PD-1 | BUV496 | 29F.1A12 | BD Biosciences | 568597 | 1:200 |
| GITR | BUV737 | DTA-1 | BD Biosciences | 741883 | 1:200 |
| GzmB | AF700 | GB11 | BD Biosciences | 560213 | 1:50 |
| Ly108 | BUV805 | 13G3 | BD Biosciences | 748564 | 1:50 |
| Anti-Puromycin | PE | 2A4 | BioLegend | 381504 | 1:100 |
| Anti-Puromycin | AF647 | 2A4 | BioLegend | 381508 | 1:100 |
| CD69 | APCCy7 | H1.2F3 | ThermoFisher | 47-0691-82 | 1:200 |
